# Comparative genomics of carbapenemase-producing *Morganella spp*

**DOI:** 10.1101/2023.03.30.534920

**Authors:** Rémy A. Bonnin, Elodie Creton, Amandine Perrin, Delphine Girlich, Cecile Emeraud, Agnès B. Jousset, Mathilde Duque, Katie Hopkins, Pierre Bogaerts, Youri Glupczynski, Niels Pfennigwerth, Marek Gniadkowski, Antoni Hendrickx, Kim van der Zwaluw, Petra Apfalter, Rainer Hartl, Vendula Heringova, Jaroslav Hrabak, Gerald Larrouy-Maumus, Eduardo P. C. Rocha, Thierry Naas, Laurent Dortet

**Affiliations:** Team “Resist” UMR1184 “Immunology of Viral, Auto-Immune, Hematological and Bacterial diseases (IMVA-HB),” INSERM, Université Paris-Saclay, CEA, LabEx LERMIT, Faculty of Medicine, Le Kremlin-Bicêtre, France; Associated French National Reference Center for Antibiotic Resistance: Carbapenemase-Producing Enterobacteriaceae, Le Kremlin-Bicêtre, France; Institut Pasteur, Université Paris Cité, CNRS UMR3525, Microbial Evolutionary Genomics, Paris 75015, France; Bacteriology-Hygiene Unit, Assistance Publique-Hôpitaux de Paris, AP-HP Paris Saclay, Bicêtre Hospital Le Kremlin-Bicêtre, France; National Institute for Health Research Health Protection Research Unit (NIHR HPRU) in Healthcare Associated Infections and Antimicrobial Resistance at Imperial College London, Hammersmith Hospital, Du Cane Road, London, Antimicrobial Resistance and Healthcare Associated Infections (AMRHAI) Reference Unit, National Infection Service, Public Health England, London, NW9 5EQ, UK; National Reference Laboratory for Monitoring of Antimicrobial Resistance in Gram-Negative Bacteria, CHU Dinant-Godinne, UCL Namur, B 5530 Yvoir, Belgium; German National Reference Centre for Multidrug-Resistant Gram-Negative Bacteria, Department of Medical Microbiology, Ruhr-University Bochum, Bochum, Germany; Department of Molecular Microbiology, National Medicines Institute, Warsaw, Poland; Laboratory for Infectious Diseases & Screening, National Institute for Public Health & the Environment (RIVM), Bilthoven, The Netherlands; National Reference Center for Antimicrobial Resistance and Nosocomial Infections, Institute for Hygiene, Microbiology and Tropical Medicine, Ordensklinikum Linz Elisabethinen, Fadingerstrasse 1, 4020 Linz, Austria; Biomedical Center, Faculty of Medicine in Pilsen, Charles University, 30100 Pilsen, Czech Republic; MRC Centre for Molecular Bacteriology and Infection, Department of Life Sciences, Faculty of Natural Sciences, Imperial College London, London, United Kingdom

**Keywords:** carbapenem, insertion sequence, antibiotic resistance, detection, colistin resistance

## Abstract

**Background:** *Morganella* are opportunistic pathogens involved in various infections. In *Morganella*, intrinsic resistance to multiple antibiotics including colistin combined with the emergence of carbapenemase-producers (CP) strongly limits the antimicrobial armamentarium.

**Methods:** From 2013 to 2021, 172 highly drug-resistant (XDR) *Morganella* isolates from 8 European countries and Canada, two reference strains from the Pasteur Institute collection and two susceptible isolates were characterized by WGS, antimicrobial susceptibility testing and biochemical tests. Complete genomes from Genbank (n=103) were included for genomic analysis. Intrinsic resistance mechanism to polymyxins was deciphered by combining genetic analysis with mass spectrometry on the lipid A.

**Findings:** *Morganella* could be separated into 4 species named *M. psychrotolerans, M. sibonii, M. morganii* and a new species represented by a unique strain. *Morganella morganii* included two subspecies: *M. morganii* subsp. *morganii* (the most prevalent) and *M. morganii* subsp. *intermedius*. Intrinsic resistance to tetracycline and conservation of metabolic pathways correlated this refined taxonomy. CP were mostly identified among five ‘high-risk’ clones of *M. morganii* subsp. *morganii*. A single nucleotide polymorphism (SNP) cut-off of 100 was used to decipher outbreaks involving this species. Cefepime-zidebactam and ceftazidime-avibactam were the most potent antimicrobials towards the 172 XDR *Morganella* spp. isolates of our collection (including 145 CP) except for metallo-β-lactamase-producers. The intrinsic resistance to polymyxins corresponds to the addition of L-Ara4N on the lipid A.

**Interpretation:** This global characterization of the widest collection of XDR *Morganella* spp. highlighted the need to clarify the taxonomy, deciphered intrinsic resistance mechanisms and paved the way for further genomic comparisons.

**Funding:** None

**RESEARCH IN CONTEXT:** *Evidence before this study:* On January 28^th^ 2022, we have searched for the terms “*Morganella*” and “carbapenemase” in all published reports available in PubMed with no language restriction. We identified a total of 43 articles and most of them (41/43) corresponded to a report of a single isolate of carbapenemase-producing *Morganella morganii*. Only one article aimed to decipher the antimicrobial susceptibility on a collection of *Proteus, Providencia* and *Morganella* isolated from global hospitalized patients with intra-abdominal and urinary tract infections. However, this collection only included 7 *M. morganii* isolates. On March 2021, when we finished the inclusions in our collection, only 104 genomes of *Morganella* spp. were available in the NCBI database. Since September 2021, very few reports were published on carbapenemase-producing Morganella with the exception of a study from Xiang G *et al*. reported 40 multi-drug resistant *M. morganii* isolates recovered from three hospitals in China from 2014 to 2020. Unfortunately, this collection included only two carbapenemase-producing *M. morganii* isolates (one OXA-48 and one IMP-1). A report of KPC-producing *M. morganii* in Japan and a longitudinal study of carbapemase-producing Enterobactrales in Taiwan that did not focused on Morganella. We also searched in PubMed for the terms ‘*Morganella sibonii*” or “*Morganella psychrotolerans*” in all published reports with no language restrictions. Our search identified a total of 20 articles. None of them was related to antimicrobial resistance and no study deciphered the *Morganella* spp. epidemiology on clinical isolates.

*Added values of this study:* This global characterization involved the widest collection of *Morganella* spp. isolates ever reported (barely doubling the number of *Morganella* spp. genomes in Genbank). In addition, 145 isolates of this worldwide collection made of 172 multidrug resistant *Morganella* spp. were carbapenemase producers for which therapeutic alternatives are scarce due to intrinsic resistance to last resort molecules, such as polymyxin. First, we found that cefepime-zidebactam and ceftazidime-avibactam were the most potent antimicrobials towards XDR *Morganella* spp. isolates except for metallo-β-lactamase-producers. Then, we observed that carbapenemase-encoding genes were present in different *Morganella* species highlighting necessary changes in the taxonomy. *Morganella* genus could be divided into 4 species named *M. psychrotolerans, M. sibonii, M. morganii* and a new species represented by a unique strain. *Morganella morganii* includes two subspecies: *M. morganii* subsp. *morganii* (the most prevalent) and *M. morganii* subsp. *intermedius*. We demonstrated that this refined taxonomy correlated with the intrinsic resistance to tetracycline, which was found only in *M. sibonii*, as well as several metabolic pathways (*e*.*g*. trehalose assimilation, type III (T3SS) and type IV secretion system (T6SS), etc.…). In addition, we highlighted five “high-risk” clones of carbapenemase-producing *M. morganii* subsp. *morganii* that have already disseminated worldwide. Combining whole genome sequencing (WGS) data with epidemiological investigations, we demonstrated that a cut-off of 100 single nucleotide polymorphisms (SNPs) could be used to discriminate clonally-related from sporadic independent isolates. This information is of the utmost importance since WGS is now considered as the reference method to identify and follow outbreaks. The intrinsic resistance of *Morganella* spp. to polymyxins was well-known but the underlying mechanism was unclear. Here, we demonstrated that the addition of L-Ara4N on the lipid A of *Morganella* is involved.

*Implications of all the available evidence:* The identification of “high-risk” clones among highly-drug resistant *Morganella* spp. paves the way of future investigations to better understand and hopefully limit the spread of these bugs. Additionally, our results identified new components and virulence factors of some *Morganella* species (e.g. T6SS and T3SS in *M. sibonii*) that deserve further investigation since they might be implicated in the bacterial lifestyle of this genus.

## INTRODUCTION

*Morganella morganii* is a facultative anaerobic Gram-negative rod belonging to Enterobacterales, firstly reported in 1907.^1^ Initially reported as *Proteus morganii*, it has been reclassified as *Morganella morganii* gen nov in 1943.^2^ Currently, *Morganella* genus is composed of two species: *M. morganii* and *Morganella psychrotolerans*.^3^ Whereas *M. morganii* is frequently encountered in clinical specimen, the psychrotolerant *M. psychrotolerans* is associated to seafood poisoning by production of histamine.^4^ In 1992, using biochemical analysis, *M. morganii* was divided into two subspecies based on the differential utilization of trehalose^5^, positive for *M. morganii* subsp. *sibonii* and negative for *M. morganii* subsp. *morganii*.

*M. morganii* is an opportunistic pathogen responsible for a wide variety of infections such as urinary tract infections, septic shock, surgical site infections, osteomyelitis, and pneumonia.^6,7^ In addition, except for a cluster of *M. morganii* infections that has been reported in the late 1970’s,^8^ no other outbreak has been studied at the microbial and genomic levels. Accordingly, the molecular epidemiology of *Morganella* spp. recovered from clinical samples has never been explored.

*Morganella* spp. isolates are intrinsically resistant to colistin, macrolides, fosfomycin, amoxicillin, first- and second-generation cephalosporins (due to the production of a class C β-lactamase, *bla*_DHA-_like) and possess decreased susceptibility to imipenem due to a low affinity of its penicillin binding protein PBP-2.^7^ The treatment of infections caused by Enterobacterales (including *Morganella*) often involves β-lactams including carbapenems. Carbapenem resistance in Enterobacterales is due to (i) the production of ESBLs or overproduced-AmpC associated with decreased outer-membrane permeability or (ii) to the production of carbapenemases, enzymes with significant hydrolytic activity towards carbapenems.^9^ The main carbapenemases are Ambler class A enzymes with mainly KPC-like enzymes, metallo-β-lactamases (MBLs) (Ambler class B) of NDM-, VIM- and IMP-type and Ambler class D carbapenem-hydrolyzing β-lactamases of OXA-48-type.^10^ Carbapenemase-producing *M. morganii* are rarely reported. However, the most prevalent carbapenemases produced by *M. morganii* are NDM-like enzymes.^11–14^ More sporadically, OXA-48- and KPC-like enzymes have also been reported in *M. morganii*.^15–18^ To our knowledge, only one report of GES-5 carbapenemase in *M. morganii* has been published.^19^ In *Morganella* spp., combining intrinsic resistance (particularly to colistin) with carbapenemase production might lead to nearly “untreatable” infections.^20^

Here, we report an extensive characterization of an international collection of carbapenemase-producing *Morganella* spp. Whole genome sequencing (WGS) combined with biochemical experiments, antimicrobial susceptibility testing and epidemiological data of several outbreaks allowed not only to decipher several mechanisms responsible for intrinsic characters of *Morganella* species, such as trehalose assimilation of *M. sibonii* and natural resistance to colistin and tetracyclin, but also to identify prevalent clonal groups inside the *Morganella* genus.

## METHODS

### Study design and strain collection

All *Morganella morganii* isolates with a reduced susceptibility profile to ertapenem or meropenem regardless imipenem susceptibility sent to the French National Reference Center (NRC) for Antimicrobial Resistance from January 1^st^ 2013 to March 1^st^ 2021 were included (n=68). The bacterial isolates referred to NRC were recovered from clinical and screening human specimens collected in french microbiology laboratories. Additionally, 104 *M. morganii* isolates suspected to produce a carbapenemase (according to EUCAST guidelines for the detection of resistance mechanisms) referred to European antimicrobial resistance reference centers were added to the collection: Germany (n=32), Belgium (n=26), England (n=17), Austria (n=3), The Netherlands (n=4), Poland (n=1), Czech Republic (n=21). One additional carbapenemase-producing *M. morganii* isolate was from Canada, two *M. morganii* subsp. *sibonii* reference strains (CIP 103648 and CIP 103649) from the Pasteur Institute collection and two susceptible isolates from Bicêtre Hospital (**Supplementary Figure S3**). Genomes of all *Morganella* spp. isolates were totally sequenced as described in supplementary methods and are deposited in Genbank database.

### Microbiology, molecular biology and bioinformatic

Bacterial identification, biochemical characterization, antimicrobial susceptibility testing and carbapenemase detection and bioinformatics methods are described in supplementary methods.

## RESULTS

### Carbapenemase-producing *Morganella* spp. in France

From January 1^st^ 2013 to March 1^st^ 2021, a total of 68 non-duplicate carbapenemase-producing *M. morganii* were collected at the French National Reference Center (NRC) among 14,672 carbapenemase-producing Enterobacterales (CPE) isolates representing 0·46% of the French CPE. These isolates were recovered from 8 different French regions (**Supplementary Figure S1B**), even though 63·2% (43/68) were recovered from the same area in South-West of France (**Supplementary Figure S1B**). Phylogenetic analysis performed on all the 68 *M. morganii* revealed that among the 53 NDM-1-producing *M. morganii*, 77·3% (41/53) belonged to the same clone recovered from 2013 to 2021 (**Supplementary Figure S1A and S1C**). This clone included 39 isolates collected in three different cities located 22 km and 72 km away in South-West of France (suggesting patient-to-patient cross-contamination), and 2 isolates collected from a distant area in North-East of France (**Supplementary Figure S1A**). The analysis of antibiotic resistance gene contents revealed that the main clone identified in South-West of France carried the *bla*_NDM-1_ carbapenemase encoding gene, the *bla*_CTX-M-15_ ESBL gene and two copies of the *bla*_DHA_-like cephalosporinase gene. The first copy corresponded to the chromosome encoded *bla*_DHA-4_ gene and the second copy was a truncated *bla*_DHA-1_ gene deleted in its 3’ end. The association of these β-lactamases encoding genes was responsible for full resistance to all β-lactams including new β-lactam-inhibitors associations (ceftazidime-avibactam, ceftolozane-tazobactam, imipenem-relebactam and meropenem-vaborbactam) (**Figure 1**). In addition, this clone I produced two 16S RNA methylases, ArmA and RmtC, and two aminoglycoside-modifying enzymes (AAC(6’)-Ib and AadA1) conferring resistance to all aminoglycosides. Resistance to quinolones was mediated by mutations in GyrA (S83I), ParC (S84I and D313E), ParE (N84K and S459Y),^21,22^ associated with the production of QnrA1. Combined with the intrinsic resistance to polymyxins and tigecycline, acquired resistance determinants were responsible to full resistance to all commonly tested molecules. Attempts to transfer the *bla*_NDM-1_ gene by conjugation or electrotransformation failed suggesting a chromosome location of the *bla*_NDM-1_ gene. Long-read sequencing of the first isolate identified in France confirmed the chromosomal location of *bla*_NDM-1_, inside a novel transposon Tn*7340* described in **Supplementary Figure S2**.

**Fig. 1.**
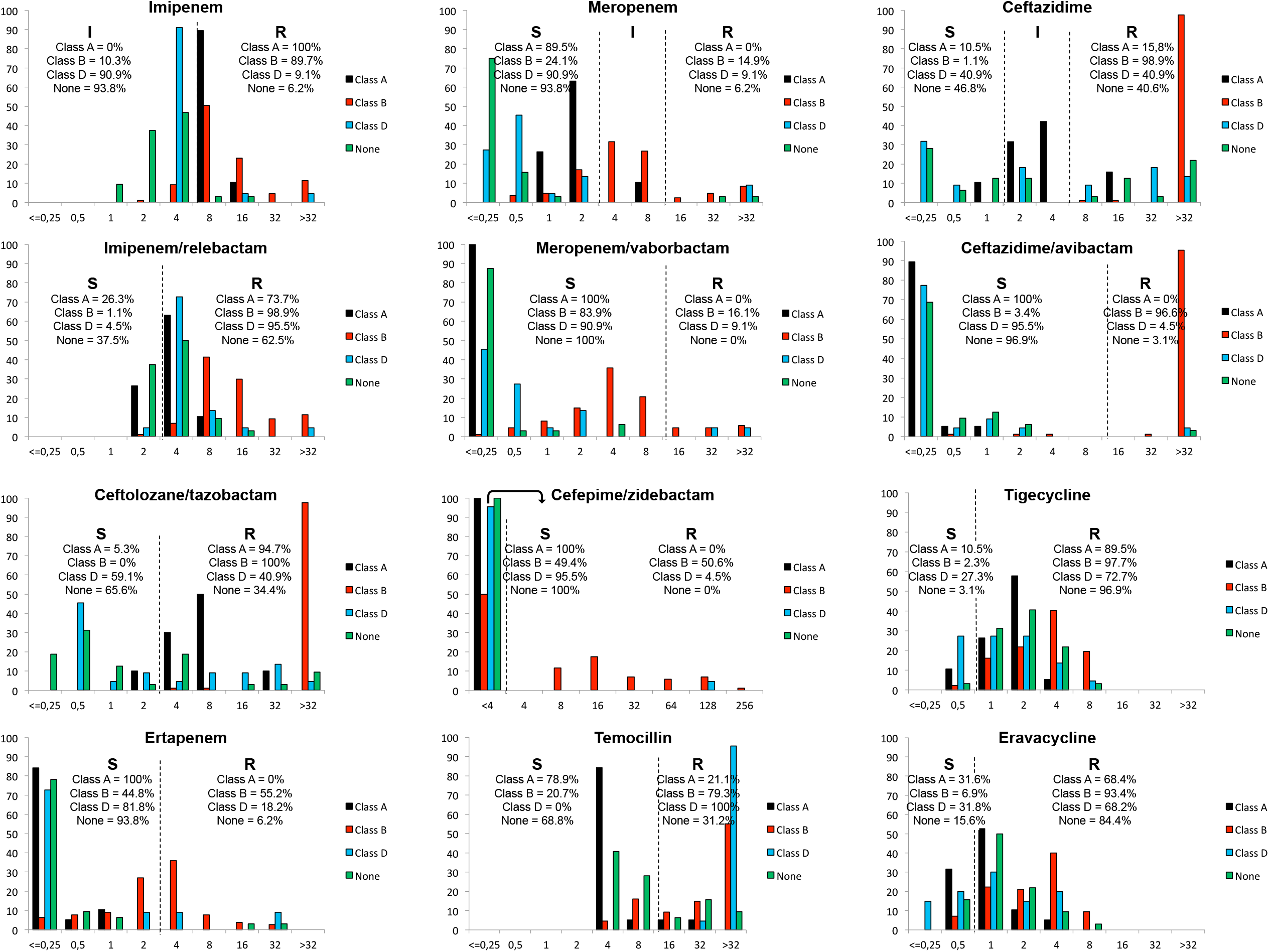

Out of this clone I, the 27 remaining isolates sent to the NRC were polyclonal and carried diverse carbapenemases (ten OXA-48, one OXA-162, twelve NDM-1, one NDM-7, one VIM-4 and one NDM-1 + VIM-1 producers) (**Supplementary Figure S1A** and **S1C**).

### Carbapenemase-producing *Morganella* spp. in Europe

To determine if the main clonal group (comprising clone I) observed in France has already spread abroad, we analyzed an additional 104 *Morganella* spp. isolates with decreased susceptibility to ertapenem or meropenem, collected from 7 reference centers across Europe from 2013 to 2021 and from Genbank (**Supplementary Figure S3**). Resistome analysis allowed the identification of a wide variety of carbapenemases produced by these isolates (**Table 1**). Of note, a wide diversity of carbapenemases was identified in Germany including NDM-1, NDM-5, VIM-1, OXA-48, OXA-181 and OXA-641 (a variant of OXA-372 reported only once in *Citrobacter freundii*).^23^ In contrast, in Czech Republic KPC-2 carbapenemase was highly prevalent (16/21) followed by OXA-48 (n=1). The resistomes of all isolates are summarized in **Supplementary Table S1**.

### Identification of “high-risk” clonal groups (CG) among carbapenemase-producing *Morganella* spp. from worldwide

First, phylogenetic analysis was conducted by creating a MASH distance similarity matrix including the whole genome of 270 *M. morganii* (68 carbapenemase-producing and 2 susceptible isolates from France, 104 from Europe, 2 from Pasteur’s Institute collection, 1 from Canada and 93 from NCBI) and 5 genomes of *M. psychrotolerans* from NCBI (**Supplementary Figure S4**). *M. psychrotolerans* was found to be very distant from *M. morganii* (less than 86% average nucleotide identity (ANI)) (**Supplementary Figure S4**). We made a phylogenetic tree SNP-calling and confirmed it using core genome maximum likelihood phylogenetic tree. Both showed that *Morganella* isolates can be roughly separated into five subpopulations (**Figure 2, Supplementary figure S4 & S5A**).

**Fig. 2.**
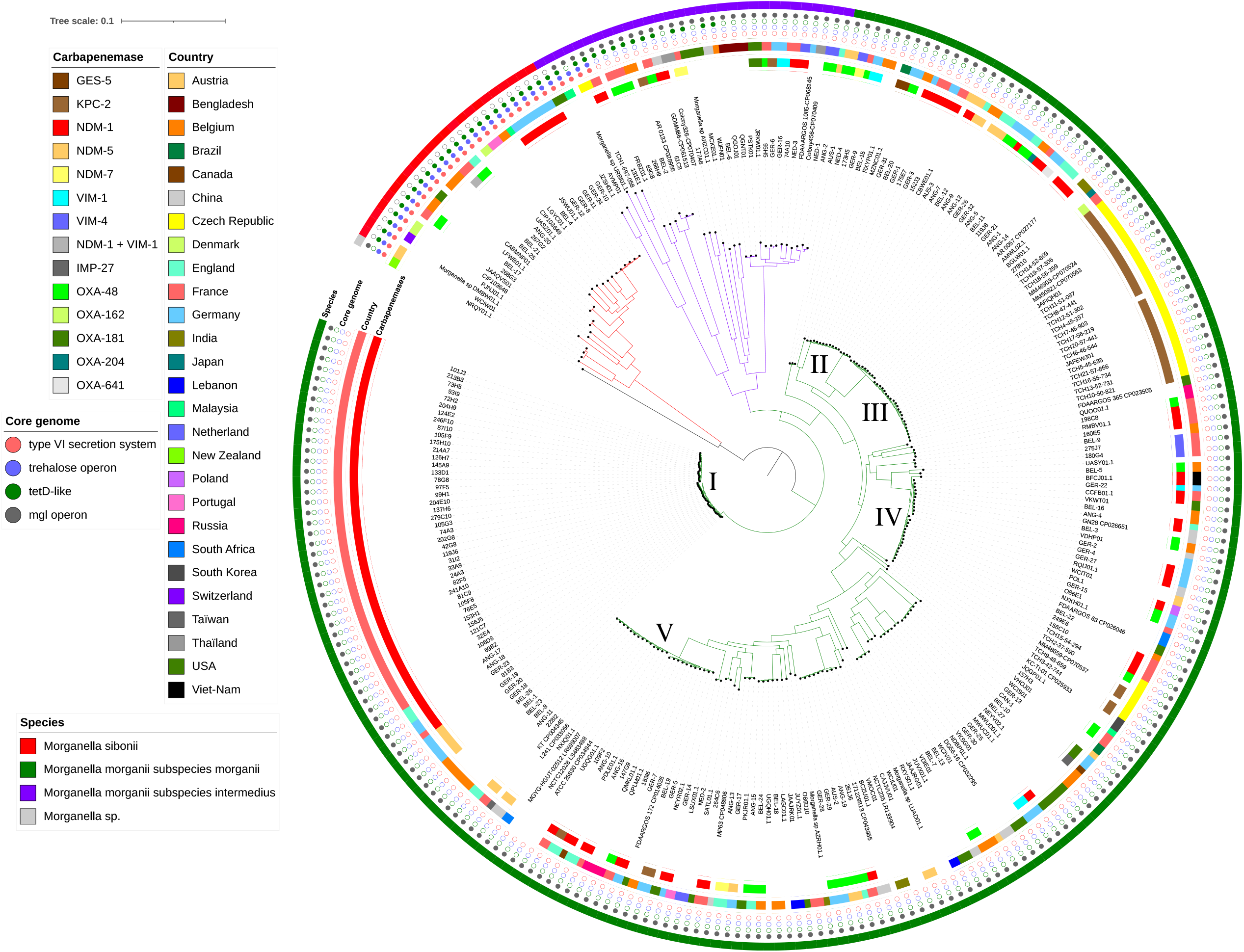

The first subpopulation corresponds to *M. sibonii* (formerly *M. morganii* subsp. *sibonii*) including 23 isolates from our study and two reference strains (CIP103648 and CIP 103649). Since, these 23 isolates possessed less than 92% identity with the *M. morganii* subsp. *morganii* and *M. psychrotolerans*, they were reclassified as an independent species named *M. sibonii* instead of a subspecies of *M. morganii*. In agreement with this new taxonomy, *M. morganii* and *M. sibonii* differed in terms of several biochemical and phenotypic characteristics (cf. below). The second and main subpopulation (n=215) corresponds to *M. morganii* subsp. *morganii*. A 3^rd^ subpopulation included 31 isolates that phylogenetically formed an independent “undefined” group with MASH distances of 94-95% with *M. morganii* subsp. *morganii* (**Figure 2** and **Supplementary Table S2**). This subpopulation was thus reclassified as a new subspecies of *M. morganii*. Since some isolates of this subspecies possessed biochemical characteristics of both *M. morganii* subsp. *morganii* and *M. sibonii* (cf. below), the name *M. morganii* subsp. *intermedius* is proposed. Finally, a unique isolate (Genbank accession number NRQY0000000) is apart from the four other populations (less than 94% ANI). This peculiar isolate was recovered from a grass grub *Costelytra sp*. in New Zealand and might be further recognized as a novel *Morganella* species if several other isolates are reported in the future.

As revealed by the phylogenetic tree, *M. morganii* subsp. *morganii*, which includes the majority of clinical isolates, might be divided in a wide diversity of CG (**Figure 2**). Among them, five CG (I to V) were more prevalent and disseminated worldwide.

The CGI, which includes the NDM-1-producing isolates of the French outbreak (n=39) also includes unrelated carbapenemase-producing isolates from Germany (n=4), United-Kingdom (n= 2), France (n=2) and Belgium (n=1). It confirms that CGI has already disseminated in Europe. A sub-tree and a SNP matrix constructed with isolates of this CGI revealed that this CG can be divided into three independent clusters (**Figure 3**). Isolates of the cluster B, which are all epidemiologically related, had less than 50 SNPs except for isolates 126H7 and 105F9 (ca. 70 to 100 SNPs). Accordingly, we decided to use a reasonable cut-off at 100 SNPs to separate clusters based on genomic results and epidemiological investigations. One French isolate 81B3 did not belong to the main French cluster (cluster B). As expected, epidemiological data revealed that the patient travelled in India and was not related to the outbreak. This strain was part of the cluster C that includes German (n=3) and English (n=2) isolates. Despite a close genetic relationship, resistomes of the isolates of this cluster C were different (**Supplementary Table S2**).

**Fig. 3.**
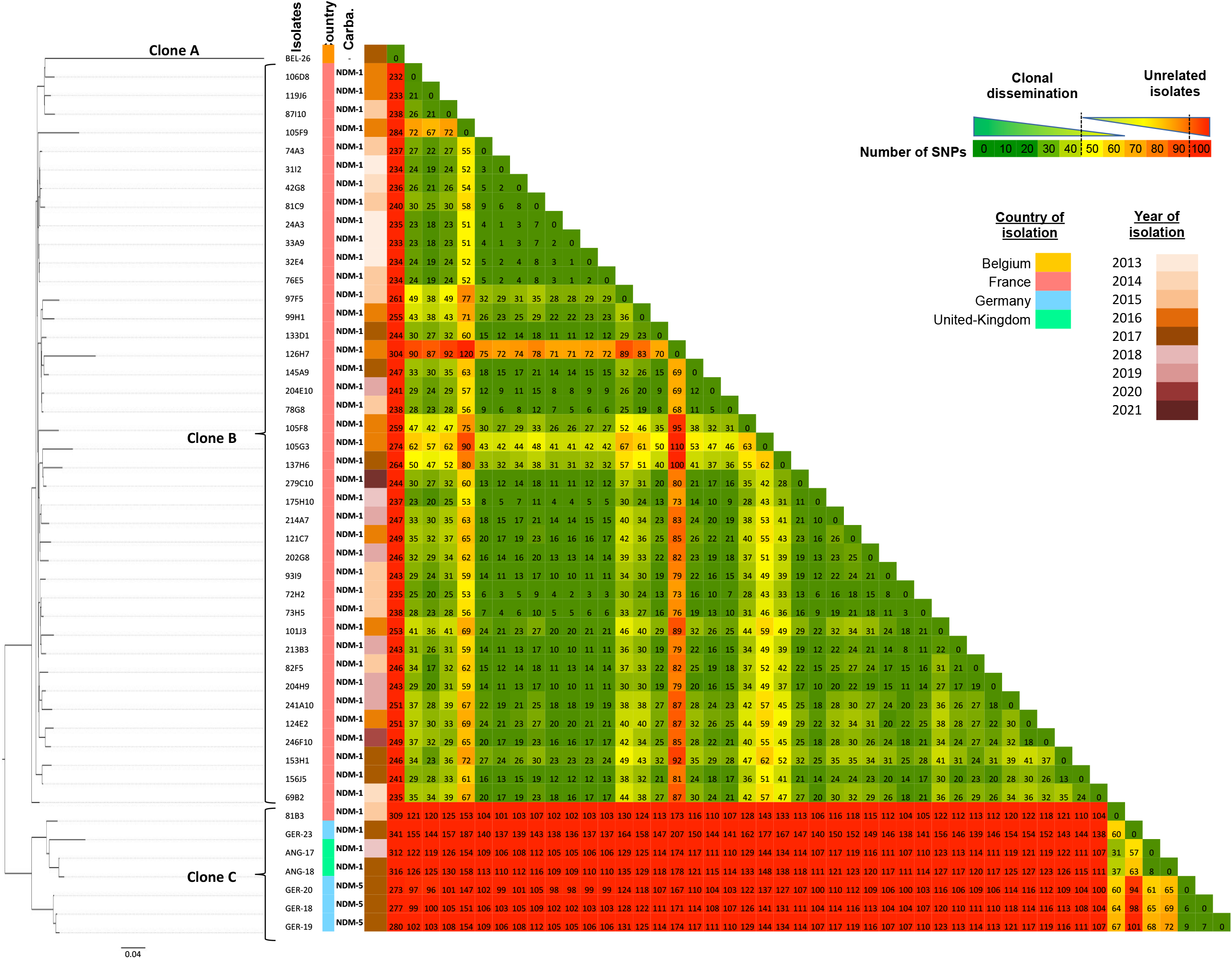

To dive deeper into the evolution of this CGI, we calculated a molecular clock based on the nucleotide diversity of the core genome using TempEST and BEAST (see Methods). The reference used was isolate 24A3, the first isolate from the French outbreak. The number of genome-wide nucleotide substitutions was correlated with isolation dates to perform a time-scale phylogeny. A root to tip regression analysis gave a R^2^ = 4.38 10^−4^ (**Supplementary Figure S5B**). The evolutionary rate was estimated at 7,782 10^-7 substitution/site/year (95% highest posterior density (HPD), 3,252.10^-7 −2,386 10^-6) soit 3,12 SNP/genome/year (1,38 −9,56). This result is in agreement with what we previously observed for *Klebsiella pneumoniae* (7.5 SNP/genome/year for ST258 & 6.2 SNP/genome/year for ST147) or *Pseudomonas aeruginosa* (7.0 SNP/genome/year).^24,25^ The last common ancestor between all strains belonging to clone I was estimated at 1972.2 and 2001 for strains belonging to the French outbreak (**Supplementary Figure S5B**).

Apart from the CGI, which was overrepresented due to the French outbreak, four additional prevalent CGs were evidenced in our collection (**Figure 2**). We analyzed the subtree of each CGs and compared the results with epidemiological data (**Supplementary Figure S6**). The CGIII is mostly composed by KPC-producers from Czech Republic that have been reported to be part of the same outbreak. Our genomic analysis confirmed epidemiological data with SNPs ranging between of 21 to 70, which confirmed the robustness of our SNP cut-off of 100. We identified several cross-country disseminations for each main CGs. As example, inside the CGIV we identified five isolates in Czech Republic carrying either OXA-48 or KPC-2 and close relationship between three isolates from France and Belgium (156C10, 249E6 and BEL-22, respectively). Out of the five main CGs, we identified three isolates from United-Kingdom (ANG-7, ANG-9 and ANG-12) with clear epidemiological link (44 to 81 SNPs) (**Supplementary Figure S7**) that were previously confirmed to be part of the same outbreak using PFGE (K. Hopkins personal data).

### Comparison of core genome from *M. morganii* and *M. sibonii*

We compared the core genomes of *M. sibonii* and *M. morganii* subsp. *morganii* (**Figure 4A**) to (i) identify unique features and (ii) to confirm results obtained with whole-genome phylogeny. Almost 2,700 genes were present in 95% of the strains both species whereas 47 genes were specific to the core genome of *M. morganii* subsp. *morganii* and 63 were specific to that of *M. sibonii* **(Figure 4A** and **Supplementary Table S3**).

**Fig. 4.**
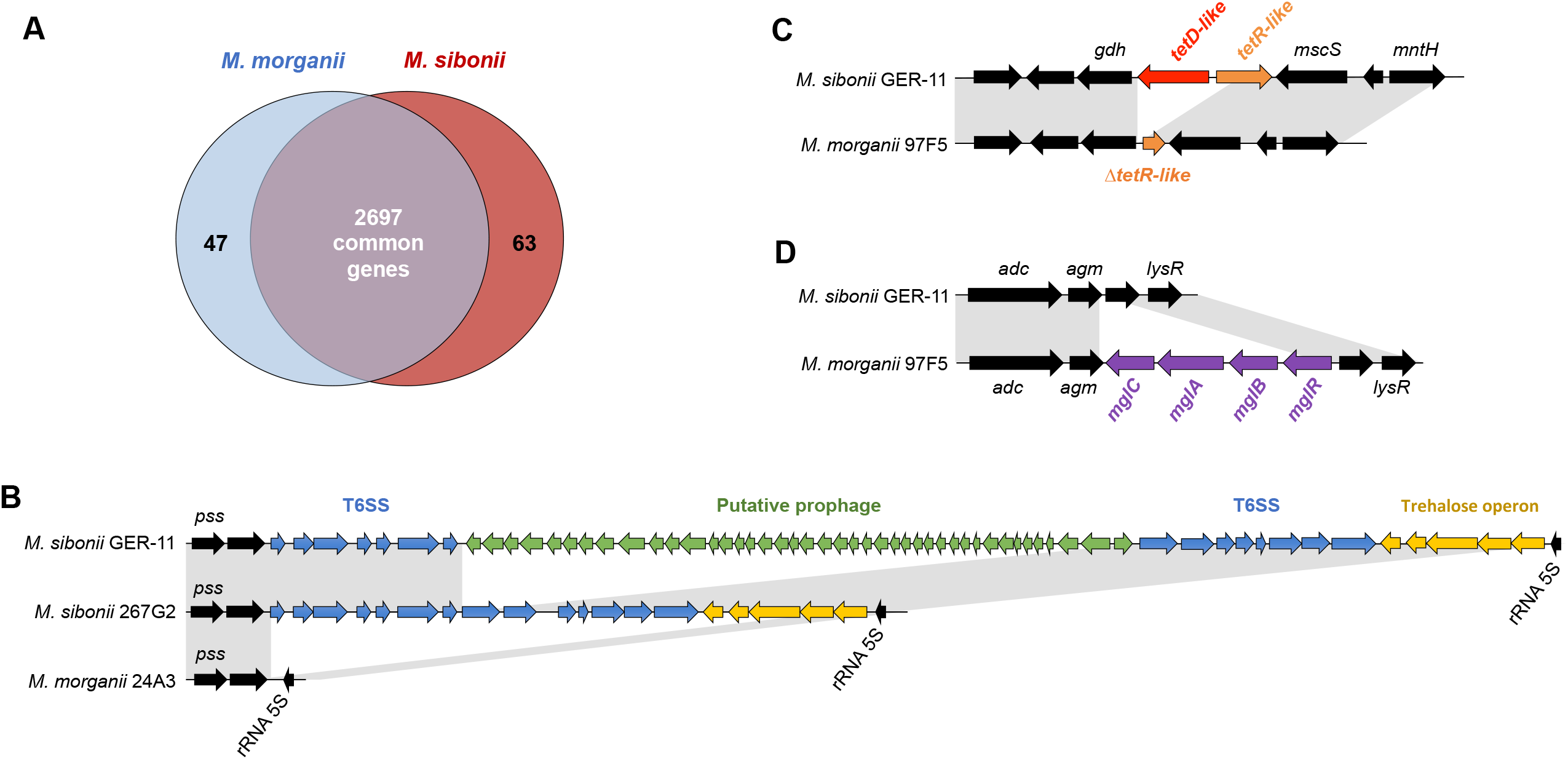

As previously described, *M. sibonii* differs from *M. morganii* subsp. *morganii* by assimilation of trehalose.^26^ The biochemical characterization using Api20E and Api50CH galleries of all isolates of our collection confirmed that all *M. sibonii* isolates were able to use trehalose as unique carbone source as opposed to *M. morganii* subsp. *morganii* strains. Among *M. sibonii* specific genes we identified the presence of an operon involved in sugar transport (**Figure 4B**). The expression of this entire operon in a *M. morganii* subsp. *morganii* strain confers the ability of this strain to utilize trehalose as sole carbon source, demonstrating the functionality of this operon. The **supplementary Figure S8** summarizes the putative role of each partner of this operon in trehalose utilization in *M. sibonii*. Immediately downstream of the trehalose operon, we systematically found a putative type VI secretion system (T6SS) (**Figure 4B**). A different putative secretion system, a type III secretion system (T3SS), is encoded in the genomes of *M. sibonii*, but not in those of *M. morganii* subsp. *morganii* (**Supplementary Table S3**).

Sometimes wrongly considered as tetracycline resistant, *M. morganii* subsp. *morganii* is intrinsically susceptible to tetracycline. Oppositely, *M. sibonii* possess in its core genome a *tetD*-like resistance gene as well as its *tetR* regulatory gene that might lead to intrinsic tetracyclin resistance (**Figure 4C**). Expressed in *E. coli*, we confirmed that the *tetD*-like gene was able to confer resistance to tetracycline (MIC > 128 mg/L) but not to tigecyline (MIC <0.25 mg/L) or eravacycline (MIC <0.25 mg/L). In *M. sibonii*, partial loss of this *tetD*-like locus in *M. morganii* subsp. *morganii* might be the result of a deletion/contraction at this locus (**Figure 4C**).

Among the genes specific to *M. morganii* subsp. *morganii* (**Table S2**), we found the whole locus *mglB/A/C* encoding the ABC transporter of galactose/methyl galactoside and its transcriptional regulator encoded by *galS*.

In line with the phylogenetic analyses (**Supplementary Figure 5A**), *M. morganii* subsp. *intermedius* has characteristics intermediate between those of *M. morganii* subsp. *morganii* and *M. sibonii*. As example, all *M. morganii* subsp. *intermedius* isolates possess the *mglB/A/C* locus specific to *M. morganii* subsp. *morganii* but few strains also possess the trehalose operon, the T6SS and the resistance gene *tetD* associated to *M. sibonii* (**Figure 2, Supplementary Table S2**). Of note, the trehalose operon was identified in *M. morganii subsp. intermedius* isolate 131E1 with a similar synteny but with only 88·2% nucleotide identity compared to the *M. sibonii* reference strain CIP103648, indicating that this operon was not recently acquired but evolved vertically within this genus.

### Antimicrobial alternatives for the treatment of highly drug resistant *Morganella* spp

The treatment of infections caused by carbapenem-resistant *Morganella* spp. is of great concern. Accordingly, we tested several last-resort antibiotics such as three carbapenems, new β-lactams-β-lactamase inhibitor associations, temocillin and two last-resort cyclins.

For each antimicrobial, MICs distributions are presented in **Figure 1** and interpreted according to EUCAST guidelines. Carbapenemase producers (n=145) were separated according to their carbapenemase class content and compared to isolates that do not produce any carbapenemase (n=32). As expected, non-carbapenemase producers showed only a moderate susceptibility to imipenem. The combination with relebactam did not significantly increase the efficiency of imipenem. Surprisingly, MIC of ertapenem remained in the susceptible range for 100% and 81·3% of Ambler class A and class D carbapenemase producers and 93·8% of non-carbapenemase producers. As expected, vaborbactam strongly restored meropenem susceptibility for all KPC producers. But this association showed a moderate effect on class D and no effect on MBL-producing isolates. The distribution of MICs of ceftazidime was heterogeneous except for MBL producers that remained highly resistant. The ceftazidime-avibactam was an accurate option for the treatment of infection caused by Ambler class A or class D carbapenemase- and non-carbapenemase-producers with 100%, 95·5% and 96·9% of susceptibility respectively. As expected, all MBL-producers were fully resistant to ceftazidime-avibactam. Temocillin showed a bi-modal distribution with 79·3% and 100% of MBL and class D carbapenemase producers being highly resistant, respectively. Oppositely, 78·9% of the Ambler class A carbapenemase producers remained susceptible. Cefepime/zidebactam demonstrated the highest efficacy with most of isolates remaining below the resistance threshold (≤ 4 mg/L). Zidebactam (formerly WCK 5222) is a β-lactamase inhibitor of the diazabicyclooctane (DBO) family and is used in combination with cefepime to potent β-lactam efficiency.^27^ In addition to its inhibition properties towards Ambler class A and class D β-lactamases, zidebactam possesses intrinsic antimicrobial activity through its binding to PBP-2 and PBP-3.^28^ However, this molecule does not show any efficacy by itself against Proteae.^27^ Accordingly, 100%, 95% and 100% of Ambler class A carbapenemase-producers, Ambler class D carbapenemase-producers and non-carbapenemase-producers were susceptible to cefepime/zidebactam, respectively. Only 49% of MBL-producing isolates were susceptible to cefepime/zidebactam. Of note, cefepime-zidebactam susceptible isolates also exhibited cefepime MIC ≤ 4 mg/L, confirming the absence of intrinsic action of zidebactam towards the PBP of *Morganella* spp. Finally, MICs of last generation cyclines (tigecycline and eravacycline) demonstrated only very moderate susceptibility to these molecules (2·3% to 31·8%).

### Deciphering polymyxin resistance mechanism in *Morganella* spp

*Morganella* spp. are known to be resistant to polymyxin at high-level. However, the underlying mechanism remained unknown. In our collection two isolates (BEL-5 and BEL-6) exhibited peculiar susceptibility to polymyxin with MICs to colistin at 2 and 0.5 mg/L respectively. To decipher the structure of the lipid A, a MALDIxin test (mass-spectrometry assay dedicated to lipid A analysis) was performed on 10 colistin-resistant isolates (MICs > 256 mg/L) (nine *M. morganii subsp. morganii* and one *M. sibonii*) and BEL-5 and BEL-6. It clearly identified a strong decrease in L-Ara4N-modified lipid A in susceptible isolates (**Supplementary Figure S9**). It demonstrated that resistance to polymyxin is caused by addition of L-Ara4N in *Morganella* genus and that some genetic events could occur, leading to the decrease in L-Ara4N modifications and acquired susceptibility to polymyxins. In *K. pneumoniae*, addition of L-Ara4N is mediated by the up-regulation of the operon *arnBCADTEF* under the control of different two-component systems (TCS) PhoP/Q and PmrA/B.^29^ By homology, nine similar TCS have been identified in this study in *Morganella* spp including PhoP/Q and QseB/C. Additionally, Guckes *et al*. demonstrated that QseB/C, involved in *quorum sensing*, could also interfere on the PmrA/B regulon.^30^ Comparative genomics were performed on BEL-5 and BEL-6 isolates. In BEL-5 isolate colistin resistance was likely due to the insertion of IS*10R* immediately upstream the *arn* operon leading to the truncation of the PmrA/Qse binding site that likely modified the expression of the *arn* operon (**Supplementary Figure S10**). In BEL-6, the PmrA/Qse binding site is intact but *arn operon* exhibited 10 times more SNPs in *arnA, arnB* & *arnC* than the whole bracketing region in comparison to the polymyxin-resistant *M. morganii* subsp. *morganii* isolate 177A6 (**Supplementary Figure S11**), suggesting an impact of these mutations in the function of the *arn* operon. However, the role of each mutation remains to be elucidated.

## DISCUSSION

The starting point of this study was to understand an outbreak of NDM-1-producing *M. morganii* subsp. *morganii* in France over a ten-year period. Surprisingly, this analysis revealed a longitudinal outbreak including isolates with very similar genetic backgrounds. The high conservation of the clone is evident from the isolates that accumulated less than 20 SNPs over a ten year-period (**Figure 3**). Using epidemiological data and SNP analyses, a cut-off value of 100 SNPs along the whole genome was considered reasonable to discriminate outbreak-related strains from non-clonally related isolates. Recently, a cut-off value of 20 SNPs along the core-genome was advocated to decipher if the *K. pneumoniae* isolates were clonally-related or not.^31^ This lower SNPs number might be explained by differences in genome comparison processes used in both studies. Indeed, in our study whole genome comparison was performed instead of a core genome comparison used for *K. pneumoniae*. We found that the molecular clock of *M. morganii* was around 5·2 SNPs/genome/year, lower than those computed with similar approaches for *K. pneumoniae* (7·5) and *P. aeruginosa* (7·0).^24,25^ However, the *in vivo* genetic diversity is very likely underestimated. Indeed, during the analysis of KPC-producing and non-KPC-producing *K. pneumoniae* ST512 within a single patient, 14 distinct genetic event was observed giving rise to a “population” of *K. pneumoniae* ST512 *in vivo* rather than a single isolate.^32^ In routine laboratory, only a single isolate is recovered for further analyses. Thus, the genetic diversity and thus the evolution rate are likely misestimated.

The comparison of carbapenemase-producing isolates from France and Europe, and its integration with the genomes in GenBank, allowed to start unravelling the population structure of *Morganella* spp. Alternative approaches, two phylogenetic reconstructions and the analysis of MASH distances, converged in showing that the *Morganella* genus should be separated into 4 species named *M. psychrotolerans, M. sibonii, M. morganii* and a new species represented by a unique strain. In addition, *Morganella morganii* could be split into two subspecies named *M. morganii* subsp. *morganii* and *M. morganii* subsp. *intermedius*. Many traits are associated with this novel taxonomic organization, including the intrinsic resistance profile to tetracycline, the conservation of metabolic pathways, or the presence of specific secretion systems.

*Morganella* spp. is intrinsically resistant to polymyxins. However, the molecular and biochemical mechanism remained unknown. In this study, the analysis of two *Morganella* isolates susceptible to colistin allowed us to demonstrate that intrinsic addition of L-Ara-4N on the lipid A *via* expression of *arnBCADTEF* leads to polymyxin resistance.

Finally, antimicrobial susceptibility testing allowed us to identify the best therapeutic options for the treatment of infections caused by MDR *Morganella* spp. As expected, we identified that relebactam do not restore imipenem susceptibility since *Morganella* spp. possess intrinsic decreased susceptibility to imipenem through PBP with low affinity to this molecule. We also demonstrated that the ceftazidime-avibactam, meropenem-vaborbactam and cefepime-zidebactam are suitable options for the treatment of Ambler class A and D carbapenemase-producing isolates as well as for non-carbapeneamse producers. Of note, as observed with Enterobacterales, the novel β-lactamase inhibitors (avibactam, relebactam, vaborbactam and zidebactam) are inefficient to restore the activity of carbapenems (imipenem or meropenem) or broad-spectrum cephalosporins (ceftazidime, cefepime) when a MBL was produced.

To conclude, this work analyzed in detail the largest ever collection of *Morganella* spp., leading to reorganization in *Morganella’s* taxonomy and identification of key specific phenotypes (trehalose assimilation, tetracycline resistance…). Our results identified new putative virulence factors in some *Morganella* species (e.g. T6SS and T3SS in *M. sibonii*) that might have implications for their lifestyle. Regarding the antimicrobial resistance potential of *Morganella* spp. (intrinsic resistance to colistin, chromosome-encoded cephalosporinase associated, acquired carbapenemase encoding genes), this genus might become a threatening issue in a next future.^20^ Accordingly, *Morganella* deserve more comprehensive studies to understand its lifestyle and its ability to acquire resistance. This should include a better knowledge of the dissemination of “high-risk clones” such as the one which was responsible for a large outbreak in France and that had already spread at least in Europe.

## ACKNOWLEDGMENTS

Transposon was named according the transposon Registry database. ^33^

