## supplementary methods for "Comparative genomics of carbapenemase-producing *Morganella spp*"

^5^ Antimicrobial Resistance and Healthcare Associated Infections (AMRHAI) Reference Unit and HCAI, Fungal, AMR, AMU and Sepsis Division, UK Health Security Agency, London, NW9 5EQ, UK

**Bacterial identification**

The bacterial identification of all 172 *Morganella* spp. collection isolates was verified by MALDI-TOF mass spectrometry (Biotyper, Bruker Daltonics).

**Biochemical characterization**

Biochemical characterization of the collection was performed using Api20E and Api50CH systems according to the manufacturer’s recommendations (BioMérieux, France).

**Antimicrobial susceptibility testing**

Antimicrobial susceptibility testing was performed by the disc diffusion method on Mueller-Hinton (MH) agar (Bio-Rad, Marnes-La-Coquette, France) and interpreted according to EUCAST guidelines as updated in 2021 (http://www.eucast.org). MICs of temocillin, cefepime/zidebactam, ceftazidime, ceftazidime/avibactam, ceftolozane/tazobactam, ertapenem, imipenem, imipenem/relabactam, meropenem, meropenem/vaborbactam, colistin, eravacycline and tigecycline were determined by broth microdilution (Sensititre^TM^ Thermofisher, France).

**Carbapenemase detection.**

Carbapenemase detection was performed using Carba NP-test as previously described, followed by an immunochromatographic detection of the carbapenemase enzyme using NG-Carba5 test (NG Biotech, Guipry, France). ^1^

**Whole genome sequencing**

All *Morganella* spp. isolates were sequenced using Illumina’s technology as previously described.^2^ *De novo* assembly and read mappings were performed using CLC Genomics Workbench v12.0 (Qiagen, Les Ulis, France). Long read sequencing was performed on *M. morganii* 97F5 using PacBio’s technologies as described previously. ^3^

**Dataset of *Morganella* spp. genomes and MASH analysis**

A dataset of 275 *Morganella* spp. genomes including 103 genomes from Genbank and 172 genomes from our collection were used for bioinformatics analysis. The similarity between genomes was estimated by calculating pairwise genetic distances with Mash v2.1.^4^ For very similar genomes, the Mash distance D strongly correlates with alignment-based measures such as the Average Nucleotide Identity (ANI) based on whole-genome sequence, with D ≈ 1 – (ANI/100). Based on current taxonomy, 3 groups were identified: *Morganella psychotolerans* (5 genomes), *Morganella morganii* subsp*. sibonii* (23 genomes) and *Morganella morganii* subsp*. morganii* (247 genomes). Only genomes from *Morganella morganii* species were kept for further analysis corresponding to 270 genomes.

**Bioinformatic analyses**

The acquired antimicrobial resistance genes were identified using Resfinder server v3.1 (<https://cge.cbs.dtu.dk/services/ResFinder/>).^5^

Sequences alignments were performed using ClustalW (<https://www.genome.jp/tools-bin/clustalw>).

The plasmids of clinical isolates were analyzed by searching replicase gene using PlasmidFinder v2.1 and by manual search for genes showing homology by blast in Genbank database with a replicase gene.

**Identification of pan- and core-genomes**

The pangenome and the core (persistent) genomes of the 270 genomes of *Morganella morganii* were identified using PanACoTA v1.2.0^6^, with default parameters (max L90 = 100 and a maximum of 999 contigs). A set of 42 genomes did not pass the quality control and was excluded. This gave our final dataset of 209 *Morganella morganii* genomes and 19 *Morganella sibonii* genomes. All 228 genomes were re-annotated with Prokka ^7^, to obtain homogeneous annotations. The pangenome of all 228 *Morganella* genomes was built*,* using the ‘pangenome’ module of PanACoTA (based on mmseqs2 v.13-45111) with default parameters.^8^ Homologous families were determined by keeping only hits with at least 80% identity and an alignment covering at least 80% of both proteins. These proteins were then clustered by single linkage. This resulted into 22,952 families of homologous proteins. Core genomes were inferred from the pangenome previously calculated, using ‘corepers’ module of PanACoTA. We built a mixed core ("persistent” in PanACoTA nomenclature) genome at 95% of the whole *Morganella morganii* species. This means that a gene is considered as core if it is present in a single copy in at least 95% of the genomes. A total of 2,697 core genes were identified. We also computed the core genomes of each of the 2 subspecies, from the *M. morganii* species pangenome. We identified a total of 2,781 core genes in the 209 *M. morganii* genomes and 2,498 genes in the 19 *M. morganii subsp. Sibonii* genomes.

**SNP-based phylogeny**

The genomes were annotated using RAST server.^9,10^ The SNPs were identified on the whole genomes using CSIphylogeny V1.4 with parameters as follow: select min depth at SNP position at 10X, minimum distance between SNPs at 10 bp, minimum SNP quality at 30. The phylogeny was then inferred using the CSIphylogeny v1.4 server ([www.cge](http://www.cge).cbs.dtu.dk/services/CSIPhylogeny/) and visualized using iTOL software v4.^11^

**Core genome phylogeny**

The core genome phylogeny was performed using the 228 genomes of *M. morganii sensu lato* and 5 *M. psychrotolerans* to confirm results obtained using whole-genome phylogeny. Trees were generated using core (persistent) genomes obtained with PanACoTA with following parameters: multiple alignments with mafft,^12^ phylogenetic tree obtained using IQ-TREE v 2.0.6^13^ with the TEST model option. A GTR+F+I+G4 model was selected according to BIC.

**Temporal signal**

Strength of temporal signal of molecular phylogeny was evaluated by root-to-tip linear regression on core-genome phylogenetic tree of clone 1 using TempEST with standard parameters v1.5.2.^14^ After the checks made with TempEST we used the resulting alignment to make a Bayesian analysis using BEAST v1.10.4. ^15^ The model used with the best fit was GTR substitution model, lognormal relaxed clock and constant population. Time estimation was performed using Tracer v1.7.2 and TreeAnnotator v2.6.0 and finally visualized using FigTree v1.4.4.^46,47^

**PCR and cloning experiments to assess trehalose assimilation and intrinsic tetracyline resistance in *Morganella sibonii***

Whole cell-DNA was extracted as previously described.^18^ DNA from *M. sibonii* GER-11 was used as template for the amplification of the trehalose operon using primers treF: 5’-ATTTGCGGTCAACACTCTCC-3’ and treR 5’-CGGCATCTGTTCTGATAACC-3’ and tetracyline resistance gene *tetD*-likeF : 5’-CGGGCAAAAACGAAAAGTCGC -3’ and *tetD*-likeR : 5’-ACGGTTCCTCTGTGTCTGAG -3’. PCR were performed using Phusion polymerase (Thermo fischer scientific, Les Ulis, France) with an annealing temperature of 55°C respectively and an extended elongation time (5 minutes) to amplify the whole trehalose operon (7,517 bp in size). These amplicons were cloned into the Zero Blunt pTOPO (KanR) cloning vector (Thermo Fisher Scientific) and then transferred into electrocomptent *E. coli* TOP10 as previously described.^2^ Plasmids were extracted from the recombinant *E. coli* using the GeneJet Plasmid miniprep kit according to the manufacturer (Thermo fischer scientific) and transferred into electrocompetent *M. morganii* O86D10. Electrocompetent *M. morganii* were prepared using the standard procedure for *E. coli* as previously described. ^18^

**MALDIxin test.**

The protocol has been performed as previously described.^19^ Briefly, a 10 μL inoculation loop of bacteria, grown on Mueller-Hinton agar for 18-24 hours, was resuspended in 200 μL of water. Mild-acid hydrolysis was performed on 100 μL of this suspension, by adding 100 μL of acetic acid 2 % v/v and incubating the mixture at 98°C for 30 min. Hydrolyzed cells were centrifuged at 17,000 *x g* for 2 min, the supernatant was discarded, and the pellet was washed 3 times with 300 μL of ultrapure water and resuspended to a density of McFarland 20 as measured using a McFarland Tube Densitometer. A volume of 0.4 μL of this suspension was loaded onto the MALDI target plate and immediately overlaid with 1.2 μL of a matrix Norharmane (Sigma-Aldrich) solubilized in chloroform/methanol 90:10 v/v to a final concentration of 10 mg/mL. For external calibration, 0.5 µL of calibration peptide was loaded along with 0.5 µL of the given calibration matrix (peptide calibration standard II, Bruker Daltonik, Germany). The samples were loaded onto a disposable MSP 96 target polished steel BC (Bruker Part-No. 8280800).

The bacterial suspension and matrix were mixed directly on the target by pipetting and the mix dried gently under a stream of air. The spectra were recorded in the linear negative-ion mode (laser intensity 95%, ion source 1 = 10.00 kV, ion source 2 = 8.98 kV, lens = 3.00 kV, detector voltage = 2652 V, pulsed ion extraction = 150 ns). Each spectrum corresponded to ion accumulation of 5,000 laser shots randomly distributed on the spot. The spectra obtained were processed with default parameters using FlexAnalysis v.3.4 software (Bruker Daltonik, Germany).

**Supplementary Figure S1. Temporal and geographical distribution of carbapenemase-producing *M. morganii sensu lato* in France.** (A) Phylogenetic tree was constructed using CSIphylogeny (cge.cbs.dtu.dk/services/CSIPhylogen) and visualized using iTOL (itol.embl.de/). (B) Geographical map of France. Each region is represented by a color. The number within region represents the number of isolates. (C) Temporal distribution of isolates by year of isolation. Colors within histogram represent the region indicated in (B).

**Supplementary Figure S2. Genetic features of the chromosome-encoded transposon Tn*7340* in NDM-1-producing *M. morganii* isolates of the Clone I.** CDS is represented by arrows. Target site duplications of IS*Ecp1*-based transposon is indicated. The genetic environment of *bla*_NDM-1_ in *M. morganii* 97F5 (A) (representative NDM-1 producing isolate of Clone I) was compared to the genome of *M. morganii* BEL-26 (B), Clone I isolate that does not contain any acquired resistance genes.

This complex structure was inserted in a transposon initially mobilized by IS*Ecp1* and also carrying the *bla*_CTX-M-15_ ESBL gene. IS*Ecp1* is known to mobilize DNA by one-ended transposition, leading to a target site duplication (TSD) of 5-bp. In this outbreak related isolate, a 5-bp TSD (TATAA) was identified surrounding the whole structure and was inserted within *yjhT* gene. Transposon Tn*7340* carried several resistance genes including the *qnrA1* quinolone-resistance gene, *smr* likely involved in antiseptic resistance, *sul1*-Δ*qacE* corresponding to a remnant of class 1 integron, the partial copy of *bla*_DHA-1_ and *bla*_NDM-1_. Inside the Tn*7340* transposon, a Tn*1548*-like composite transposon has inserted upstream of *bla*_NDM-1_. It is flanked by two copies of IS*26* and is responsible for the spread of 16S rRNA methylase *armA* gene. Finally, this transposon also carried two aminoglycoside acetyltransferases *aac(6’)-Ib* and *aac(6’)-Ib-cr* (which is additionally responsible for quinolone resistance), sulfamide and antiseptic resistance genes *sul1*-Δ*qacE,* and macrolide resistance genes *msrE and mphE*.

**Supplementary Figure S3. Distribution map**

**Supplementary Figure S4. MASH distance analysis.** Genomes similarity were estimated by calculating pairwise genetic distances with Mash v2.1.^4^

**Supplementary Figure S5. A. Phylogenetic tree of the clade, using the core genome maximum likelihood phylogeny. B. Molecular clock evolution of *M. morganii* subsp. *morganii* isolates of the clone I**.

**Supplementary Figure S6.** **Analysis of SNPs** **in *M. morganii* subsp. *Morganii* clone II, clone III, clone IV and clone V.** The phylogenetic tree were obtained by comparing isolates from *M. morganii* subsp. *morganii* belonging to the same clone. The matrix represents the SNPs between genomes. The Year of isolation, Country and carbapenemase of each isolate is indicated at the vicinity of the isolate name.

**Supplementary Figure S7.** **Analysis of SNPs in *M. morganii* subsp. *morganii* epidemiologically linked by PFGE from United-Kingdom.** The phylogenetic tree was obtained by comparing isolates ANG-12, ANG-7, ANG-9 and BEL-12. A SNP-based matrix was visualized. Year of isolation, Country and carbapenemase of each isolate is indicated at the vicinity of the isolate name

**Supplementary Figure S8. Role of each partner in trehalose utilization in *Morganella sibonii* as compared to *Escherichia coli*.**

The first gene is a transcriptional regulator likely involved in expression of the operon when trehalose is present as demonstrated previously. The second gene *(treB*) encodes the PTS that phosphorylates trehalose in trehalose-6P. The most interesting feature with this operon is the presence of *trePP* that encodes the trehalose-6-phosphate phosphorylase. This enzyme is able to cleave trehalose-6P in glucose-6P and glucose-1P. It has been firstly identified in *Lactococcus lactis.* **Indeed, Andersson *et al.* showed the catalysis of trehalose-6P in *L. lactis* and *Enterococcus faecalis* and evidenced the role of the** β-phosphoglucomutase encoded in the homologous operon in *L. lactis.* However, *L. lactis* is a gram-positive bacterium and thus does not possess outer-membrane. In *E. coli*, the uptake of trehalose starts by the passage through the maltoporin LamB. The entry of trehalose in the periplasm of *M. sibonii* remain to be elucidated but the operon organization suggest that the last gene of this operon encoding a putative glycoporin might play a role there.

Of note, no mobile element was identified around this operon indicating its acquisition. Therefore, it suggested that this operon had probably been lost *M. morganii* subsp. *morganii* during its evolution. Another indication of this lost is the presence of this operon in *M. morganii* subsp. *intermedius* subspecies that possess characters of both *M. morganii* subsp. *morganii* and *M. siboni* (**Figure 3**). The analysis of nucleotide polymorphism showed that this locus could have evolved in the same manner in *M. sibonii* and in *M. morganii* subsp. *intermedius*, indicating that no recent genetic transfer should have occurred between these species.

**Supplementary Figure S9. Representative mass spectra of native and modified lipid A of *Morganella* spp*. isolates* acquired using the linear negative-ion mode of a matrix-assisted laser desorption ionization (MALDI) Biotyper Sirius system (Bruker Daltonics).** Native lipid A are indicated in green (*m/z* 1796 and *m/z* 2033), L-Ara4N modified lipid A are indicated in red (*m/z* 1929 and *m/z* 2166). The structures of native and modified lipid A are shown on top of their respective peaks. Colistin MICs are indicated for each isolates.

**Supplementary Figure S10. Schematic representation of the region encoding *arn* operon in the colistin resistant (wild-type) *M. morganii* subsp. *morganii* 97F5 isolate and colistin susceptible *M. morganii* subsp. *morganii* BEL-5 isolate.** PmrA/QseB consensus binding-site is indicated in the upper part of the figure. PmrA/QseB binding-site is underlined and conserved nucleotide is indicated in red. Initiation codon is indicated in bold. Inverted repeat sequences is indicated in bold and italicized.

**Supplementary Figure S11. Mapping of the reads of the colistin susceptible *M. morganii* subsp. *morganii* BEL-6 on the *arn* region of the colistin resistant (wild-type) *M. morganii* subsp. *morganii* 177A6 isolate**. The *arn* operon is represented by purple arrows. Other genes are indicated by yellow arrows. SNPs are indicated by vertical lines.

**Supplementary Table S1.** Characteristics of the 275 genomes of *Morganella* spp. analyzed in this study

Cf (Excel file)

**Supplementary Table S2.** Specific genes identified in the main clones of *M. morganii* subsp. *morganii*

| family number | gembase name | gene_name | annotation |
| --- | --- | --- | --- |
| Clone I |  |  |  |
| 246 | MOMO.2103.00072.0005i_01318 | NA | \| hypothetical protein \| NA \| NA \| NA AntA/AntB family antirepressor |
| 310 | MOMO.2103.00072.0005i_01317 | NA | \| hypothetical protein \| NA \| NA \| NA |
| 317 | MOMO.2103.00072.0016i_02831 | NA | \| hypothetical protein \| NA \| NA \| NA DNA primase (Morganella phage IME1369_03) |
| 815 | MOMO.2103.00072.0022b_03227 | NA | \| hypothetical protein \| NA \| NA \| NA Elp3 domain-containing protein |
| 5027 | MOMO.2103.00072.0005i_01314 | NA | \| hypothetical protein \| NA \| NA \| NA |
| 5051 | MOMO.2103.00072.0012b_02370 | NA | \| hypothetical protein \| NA \| NA \| NA DUF262 domain-containing protein |
| 5340 | MOMO.2103.00072.0003i_00689 | NA | \| hypothetical protein \| NA \| NA \| NA Glycosyl hydrolases |
| 5646 | MOMO.2103.00072.0016i_02810 | NA | \| putative multidrug-efflux transporter \| NA \| similar to AA sequence:UniProtKB:P9WJX3 \| NA |
| 11406 | MOMO.2103.00072.0003i_00660 | NA | \| hypothetical protein \| NA \| NA \| NA Putative Colicin-E6 (Ribonuclease) |
| 11626 | MOMO.2103.00072.0016i_02811 | NA | \| hypothetical protein \| NA \| NA \| NA |
| 12422 | MOMO.2103.00072.0005i_01315 | intS_2 | \| Prophage integrase IntS \| NA \| similar to AA sequence:UniProtKB:P37326 \| COG:COG0582 |
| 12496 | MOMO.2103.00072.0002i_00325 | NA | \| hypothetical protein \| NA \| NA \| NA Ferrichrome-iron receptor |
| 19419 | MOMO.2103.00072.0004i_00919 | NA | \| hypothetical protein \| NA \| NA \| NA |
| Clone II |  |  |  |
| 5062 | MOMO.2103.00062.0006i_01218 | fatA_1 | \| Ferric-anguibactin receptor FatA \| NA \| TonB **Ferrichrome-iron receptor** |
| 5610 | MOMO.2103.00062.0018i_02648 | NA | \| hypothetical protein \| NA \| NA \| Pribosyltran domain-containing protein |
| 6062 | MOMO.2103.00062.0038i_03789 | NA | \| hypothetical protein \| NA \| NA \| NA |
| 8180 | MOMO.2103.00062.0018i_02645 | NA | \| hypothetical protein \| NA \| NA \| NA |
| 8405 | MOMO.2103.00062.0018i_02637 | NA | \| hypothetical protein \| NA \| NA \| NA |
| 8625 | MOMO.2103.00062.0018i_02636 | NA | \| hypothetical protein \| NA \| NA \| putative transposase |
| 12003 | MOMO.2103.00062.0018i_02652 | NA | \| hypothetical protein \| NA \| NA \| NA |
| 12434 | MOMO.2103.00062.0018i_02650 | NA | \| hypothetical protein \| NA \| NA \| NA |
| 12786 | MOMO.2103.00062.0018i_02649 | NA | \| hypothetical protein \| NA \| NA \| putative TIGR03747 family integrating conjugative element membrane protein |
| 19170 | MOMO.2103.00062.0006i_01219 | fatA_2 | \| Ferric-anguibactin receptor FatA \| NA \| TonB **Ferrichrome-iron receptor** |
| 19666 | MOMO.2103.00062.0018i_02647 | NA | \| hypothetical protein \| NA \| NA \| NA |
| 22584 | MOMO.2103.00062.0011i_01694 | NA | \| hypothetical protein \| NA \| NA \| NA |
| 22636 | MOMO.2103.00062.0018i_02651 | NA | \| hypothetical protein \| NA \| NA \| NA |
| Clone III |  |  |  |
| None |  |  |  |
| Clone IV |  |  |  |
| None |  |  |  |
| Clone V |  |  |  |
| 12840 | MOMO.2103.00005.0001i_01598 | NA | \| hypothetical protein \| NA \| NA \| **DUF4431 domain-containing protein** |

**Supplementary Table S3. Unique genes present in *Morganella morganii* subsp. *morganii* and *Morganella sibonii*.**

| Annotations | Gene names | Putative functions |
| --- | --- | --- |
| ***Morganella morganii* subsp. *morganii*** | | |
| MOMO.2103.00001.0001i_00107 |  | hypothetical protein |
| MOMO.2103.00001.0001i_00132 | fimA_1 | Type-1 fimbrial protein, A chain |
| MOMO.2103.00001.0001i_00688 |  | hypothetical protein |
| MOMO.2103.00001.0001i_00915 |  | hypothetical protein |
| MOMO.2103.00001.0001i_00940 | cynR | HTH-type transcriptiol regulator CynR |
| MOMO.2103.00001.0001i_00983 | mglA_2 | Galactose/methyl galactoside import ATP-binding protein MglA 7.5.2.11 |
| MOMO.2103.00001.0001i_00984 | mglB | D-galactose-binding periplasmic protein |
| MOMO.2103.00001.0001i_00985 | galS | HTH-type transcriptiol regulator GalS |
| MOMO.2103.00001.0001i_01082 |  | hypothetical protein |
| MOMO.2103.00001.0001i_01083 |  | hypothetical protein |
| MOMO.2103.00001.0001i_01084 | exsA_1 | Exoenzyme S synthesis regulatory protein ExsA |
| MOMO.2103.00001.0001i_01085 |  | Putative binding protein |
| MOMO.2103.00001.0001i_01086 | cysW_1 | Sulfate transport system permease protein CysW |
| MOMO.2103.00001.0001i_01087 | fbpC | Fe(3+) ions import ATP-binding protein FbpC 7.2.2.7 |
| MOMO.2103.00001.0001i_01116 |  | hypothetical protein |
| MOMO.2103.00001.0001i_01120 |  | hypothetical protein |
| MOMO.2103.00001.0001i_01307 | csy1 | CRISPR-associated protein Csy1 |
| MOMO.2103.00001.0001i_01308 | csy2 | CRISPR-associated protein Csy2 |
| MOMO.2103.00001.0001i_01311 |  | hypothetical protein |
| MOMO.2103.00001.0001i_01453 | lpfA | putative major fimbrial subunit LpfA |
| MOMO.2103.00001.0001i_01454 | focC | Chaperone protein FocC |
| MOMO.2103.00001.0001i_01456 |  | hypothetical protein |
| MOMO.2103.00001.0001i_01560 |  | hypothetical protein |
| MOMO.2103.00001.0001i_01612 |  | hypothetical protein |
| MOMO.2103.00001.0001i_01656 | infC | Translation initiation factor IF-3 |
| MOMO.2103.00001.0001i_01777 |  | hypothetical protein |
| MOMO.2103.00001.0001i_01905 |  | hypothetical protein |
| MOMO.2103.00001.0001i_01941 | spvC | MAPK phosphothreonine lyase 4.2.3.- |
| MOMO.2103.00001.0001i_01947 |  | hypothetical protein |
| MOMO.2103.00001.0001i_01969 |  | hypothetical protein |
| MOMO.2103.00001.0001i_01989 |  | hypothetical protein |
| MOMO.2103.00001.0001i_02056 |  | hypothetical protein |
| MOMO.2103.00001.0001i_02366 |  | hypothetical protein |
| MOMO.2103.00001.0001i_02369 |  | hypothetical protein |
| MOMO.2103.00001.0001i_02386 | pgrR_1 | HTH-type transcriptiol regulator PgrR |
| MOMO.2103.00001.0001i_02416 |  | hypothetical protein |
| MOMO.2103.00001.0001i_02567 |  | hypothetical protein |
| MOMO.2103.00001.0001i_02673 | cspD | Cold shock-like protein CspD |
| MOMO.2103.00001.0001i_02708 | aarA | Rhomboid protease AarA 3.4.21.105 |
| MOMO.2103.00001.0001i_02814 |  | hypothetical protein |
| MOMO.2103.00001.0001i_02987 | xapB | Xanthosine permease |
| MOMO.2103.00001.0001i_02988 | pu | Purine nucleoside phosphorylase 1 2.4.2.1 |
| MOMO.2103.00001.0001i_02989 | gC_2 | N-acetylglucosamine repressor |
| MOMO.2103.00001.0001i_03107 |  | hypothetical protein |
| MOMO.2103.00001.0001i_03285 |  | hypothetical protein |
| MOMO.2103.00001.0001i_03338 |  | hypothetical protein |
| MOMO.2103.00001.0001i_03383 |  | hypothetical protein |
| ***Morganella* *sibonii*** | | |
| MOSI.2103.00001.0001i_00097 | irtB | x Iron import ATP-binding/permease protein IrtB |
| MOSI.2103.00001.0001i_00100 |  | hypothetical protein |
| MOSI.2103.00001.0001i_00205 |  | hypothetical protein |
| MOSI.2103.00001.0001i_00664 |  | hypothetical protein |
| MOSI.2103.00001.0001i_00699 | smc_1 | Chromosome partition protein Smc |
| MOSI.2103.00001.0001i_00704 |  | hypothetical protein |
| MOSI.2103.00001.0001i_00913 | treR | HTH-type transcriptiol regulator TreR |
| MOSI.2103.00001.0001i_00914 | treB | PTS system trehalose-specific EIIBC component |
| MOSI.2103.00001.0001i_00915 | trePP | Trehalose 6-phosphate phosphorylase 2.4.1.216 |
| MOSI.2103.00001.0001i_00916 | yvdM | Beta-phosphoglucomutase 5.4.2.6 |
| MOSI.2103.00001.0001i_00917 |  | Putative glycoporin |
| MOSI.2103.00001.0001i_00918 |  | T6SS component TssM |
| MOSI.2103.00001.0001i_00919 |  | T6SS component ImpK |
| MOSI.2103.00001.0001i_00920 |  | T6SS component ImpJ |
| MOSI.2103.00001.0001i_00921 |  | hypothetical protein |
| MOSI.2103.00001.0001i_00922 |  | hypothetical protein |
| MOSI.2103.00001.0001i_00923 |  | hypothetical protein |
| MOSI.2103.00001.0001i_00926 |  | hypothetical protein |
| MOSI.2103.00001.0001i_00927 |  | VgrG related protein |
| MOSI.2103.00001.0001i_00929 |  | hypothetical protein |
| MOSI.2103.00001.0001i_00930 |  | T6SS component ImpG |
| MOSI.2103.00001.0001i_00931 | hcp1_1 | Protein hcp1 |
| MOSI.2103.00001.0001i_00932 |  | T6SS component ImpC |
| MOSI.2103.00001.0001i_00933 |  | T6SS component ImpB |
| MOSI.2103.00001.0001i_00934 |  | hypothetical protein |
| MOSI.2103.00001.0001i_01002 |  | hypothetical protein |
| MOSI.2103.00001.0001i_01163 |  | hypothetical protein |
| MOSI.2103.00001.0001i_01283 |  | hypothetical protein |
| MOSI.2103.00001.0001i_01371 |  | hypothetical protein |
| MOSI.2103.00001.0001i_01453 |  | hypothetical protein |
| MOSI.2103.00001.0001i_01520 | pgrR_1 | HTH-type transcriptiol regulator PgrR |
| MOSI.2103.00001.0001i_01955 |  | hypothetical protein |
| MOSI.2103.00001.0001i_01956 |  | hypothetical protein |
| MOSI.2103.00001.0001i_02091 |  | hypothetical protein |
| MOSI.2103.00001.0001i_02229 | aadK | Aminoglycoside 6-adenylyltransferase 2.7.7.- |
| MOSI.2103.00001.0001i_02396 | infC | Translation initiation factor IF-3 |
| MOSI.2103.00001.0001i_02449 |  | hypothetical protein |
| MOSI.2103.00001.0001i_02509 |  | hypothetical protein |
| MOSI.2103.00001.0001i_02526 |  | hypothetical protein |
| MOSI.2103.00001.0001i_02624 |  | hypothetical protein |
| MOSI.2103.00001.0001i_02626 | focC_2 | Chaperone protein FocC |
| MOSI.2103.00001.0001i_02798 |  | hypothetical protein |
| MOSI.2103.00001.0001i_02828 |  | hypothetical protein |
| MOSI.2103.00001.0001i_02830 | yopB | Protein YopB |
| MOSI.2103.00001.0001i_02831 |  | hypothetical protein |
| MOSI.2103.00001.0001i_02833 | invA_1 | Invasion protein InvA |
| MOSI.2103.00001.0001i_02834 |  | hypothetical protein |
| MOSI.2103.00001.0001i_02835 |  | hypothetical protein |
| MOSI.2103.00001.0001i_02836 | yscN | putative ATP synthase YscN 7.1.2.2 |
| MOSI.2103.00001.0001i_02837 |  | hypothetical protein |
| MOSI.2103.00001.0001i_02839 | yscQ | Yop proteins translocation protein Q |
| MOSI.2103.00001.0001i_02840 | spaP_1 | Surface presentation of antigens protein SpaP |
| MOSI.2103.00001.0001i_02841 | spaQ_1 | Surface presentation of antigens protein SpaQ |
| MOSI.2103.00001.0001i_02842 |  | T3SS inner membran protein |
| MOSI.2103.00001.0001i_02843 | yscU | Yop proteins translocation protein U |
| MOSI.2103.00001.0001i_02911 |  | hypothetical protein |
| MOSI.2103.00001.0001i_03038 |  | hypothetical protein |
| MOSI.2103.00001.0001i_03039 | cynR | HTH-type transcriptiol regulator CynR |
| MOSI.2103.00001.0001i_03041 |  | hypothetical protein |
| MOSI.2103.00001.0001i_03042 |  | hypothetical protein |
| MOSI.2103.00001.0001i_03324 | gtf3 | Glucosyltransferase 3 2.4.1.- |
| MOSI.2103.00001.0001i_03325 |  | hypothetical protein |
| MOSI.2103.00001.0001i_03979 |  | hypothetical protein |
