## Supplementary figures and images for "Comparative genomics of carbapenemase-producing *Morganella spp*"

### Supplementary Figure S1

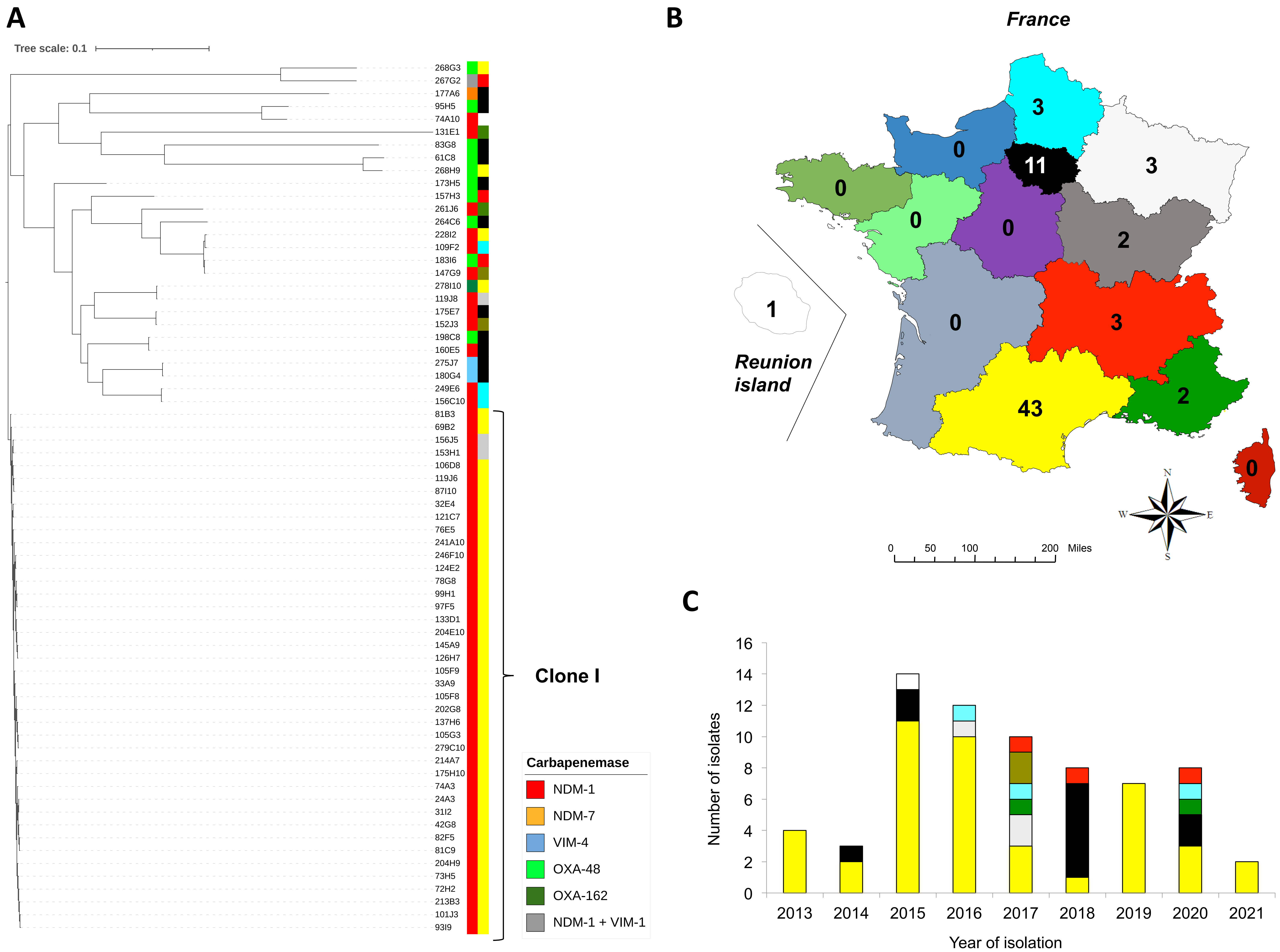

### Supplementary Figure S2

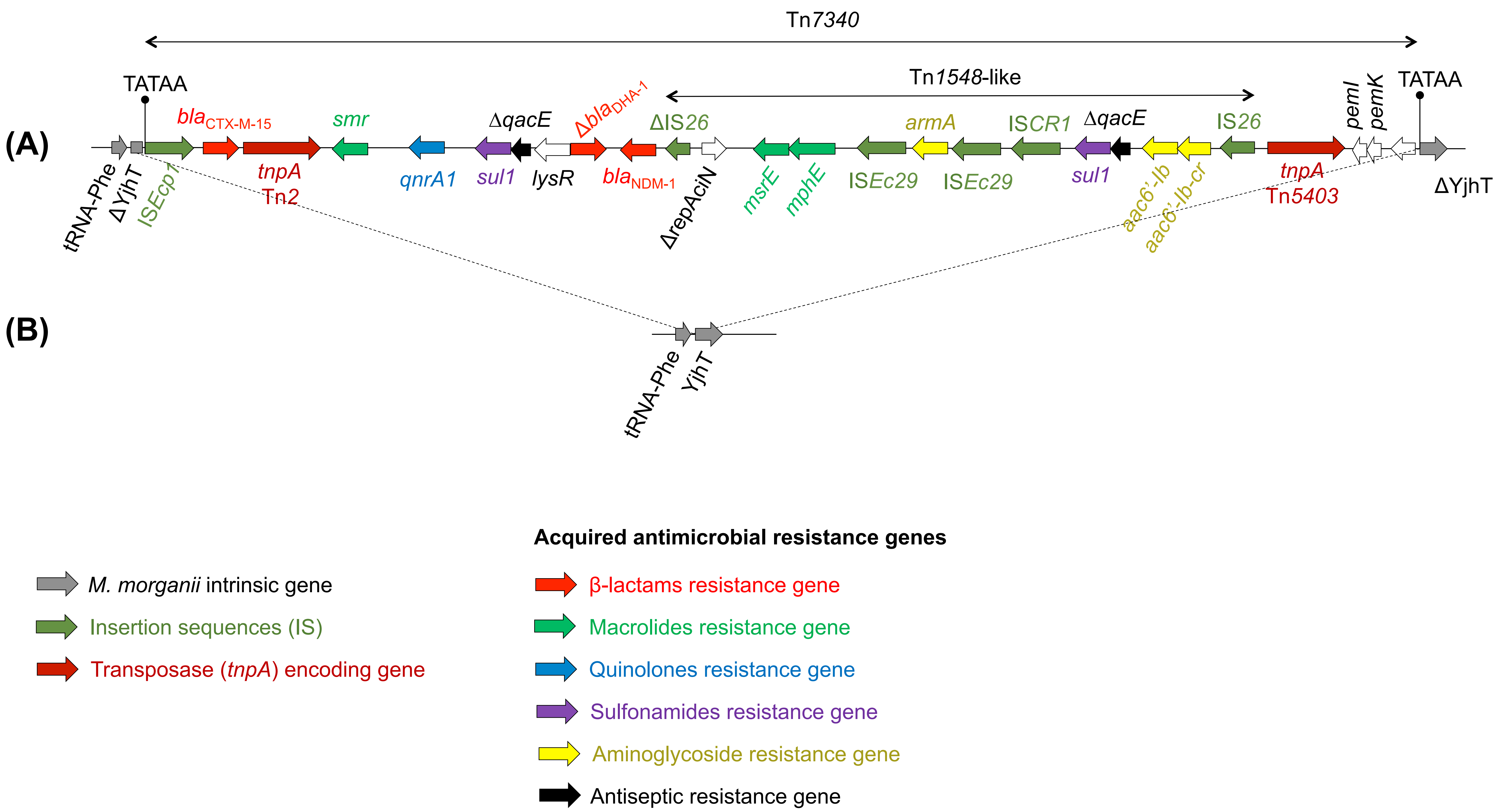

### Supplementary Figure S3

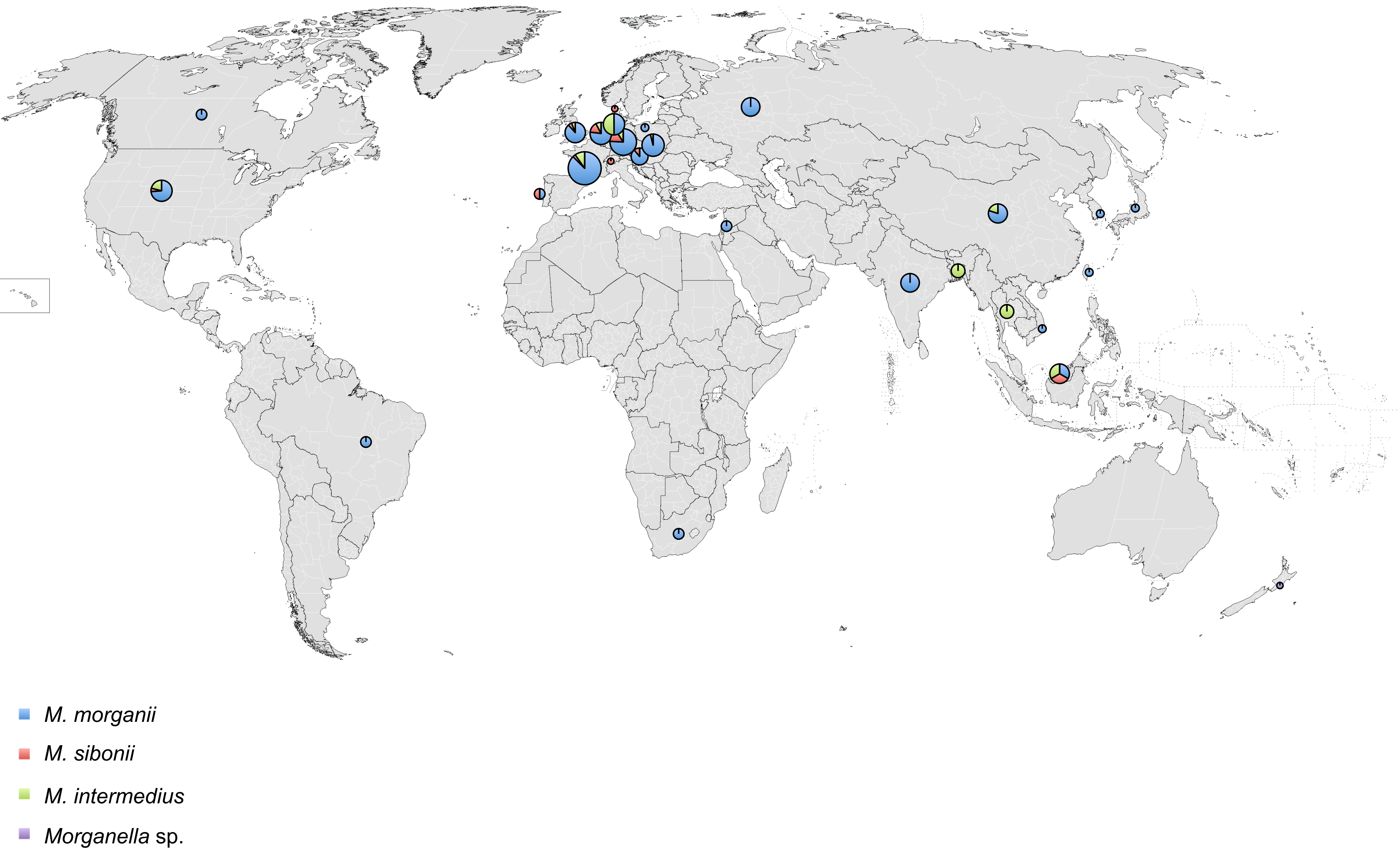

### Supplementary Figure S4

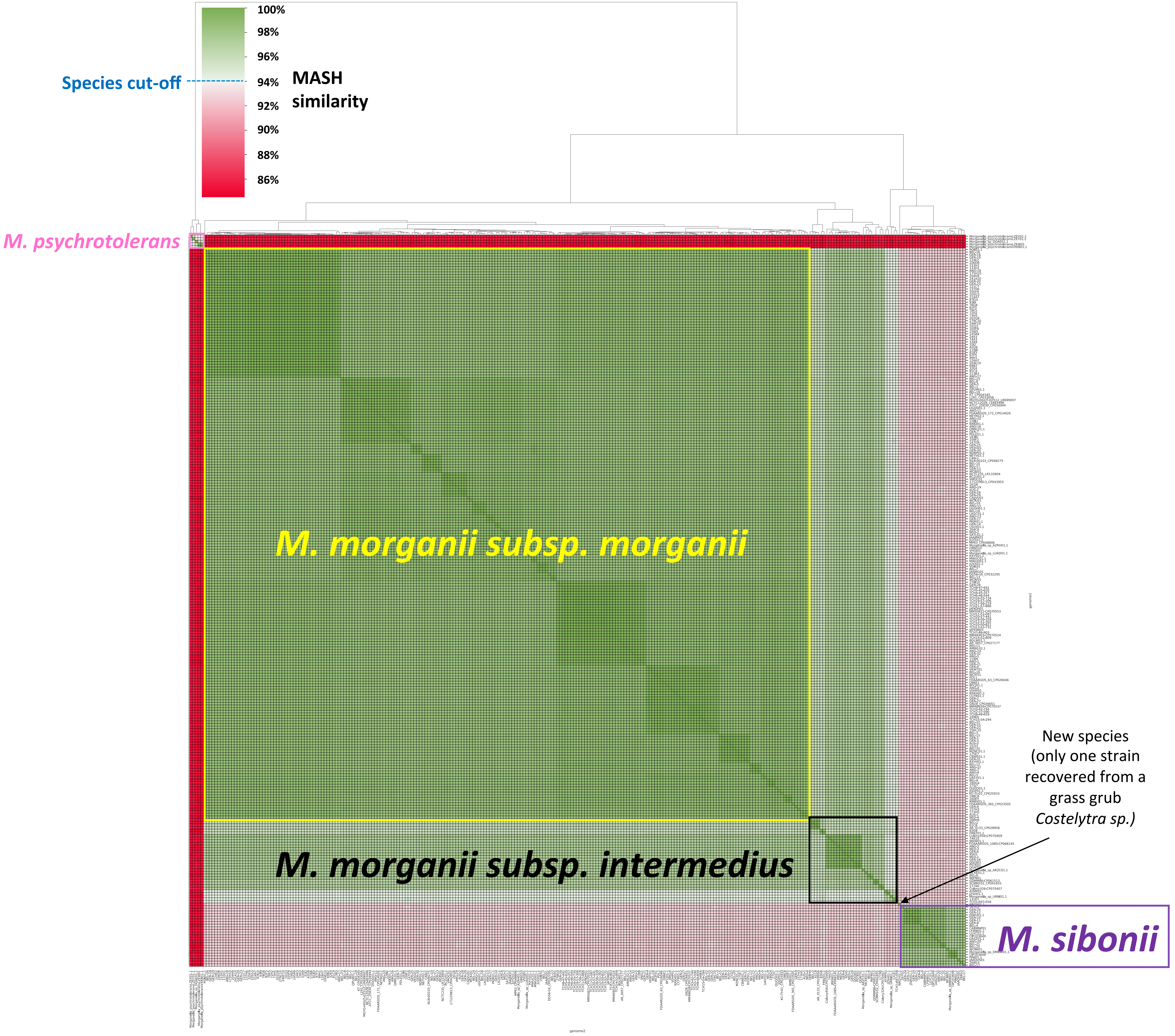

### Supplementary Figure S5

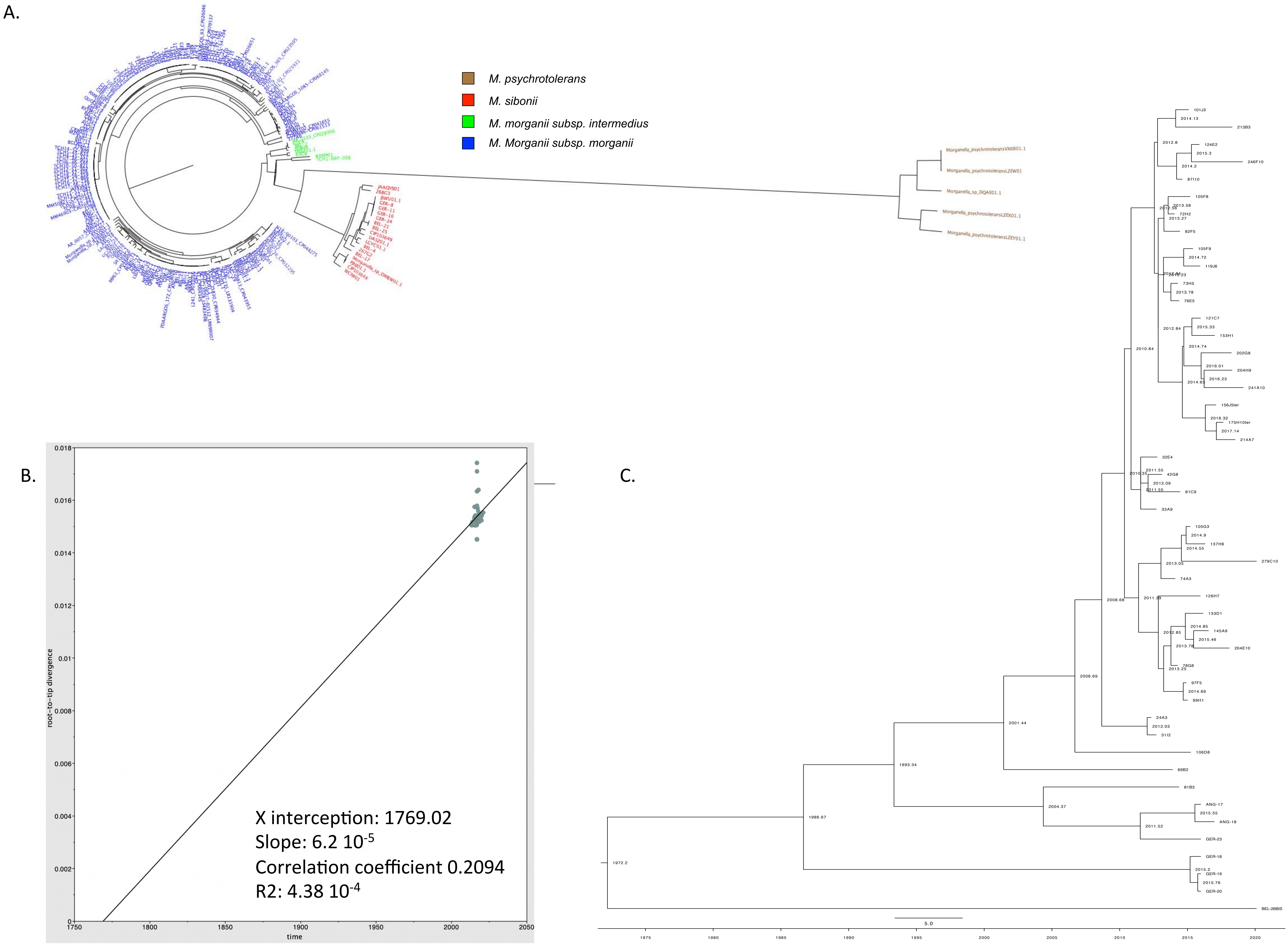

### Supplementary Figure S7

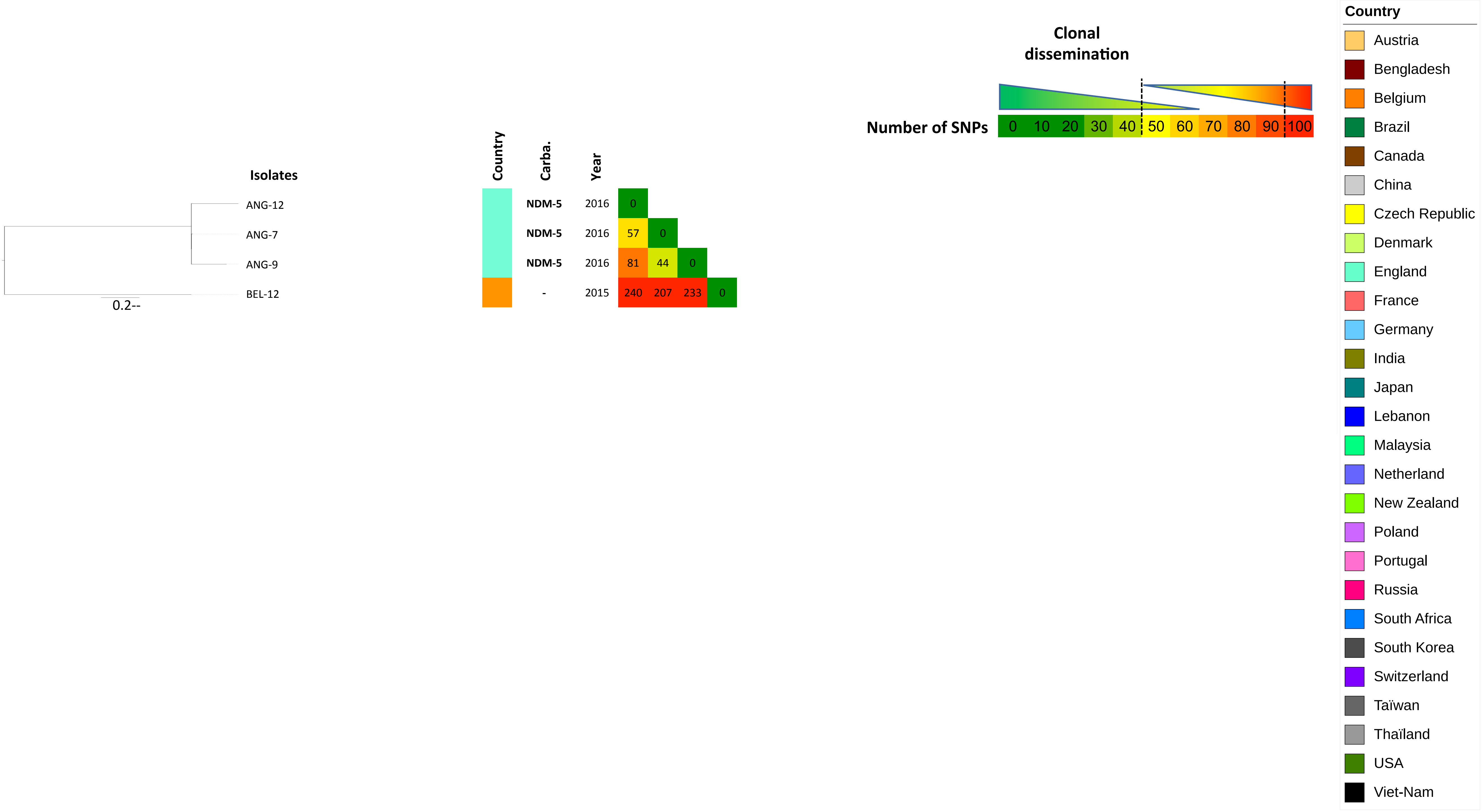

### Supplementary Figure S8

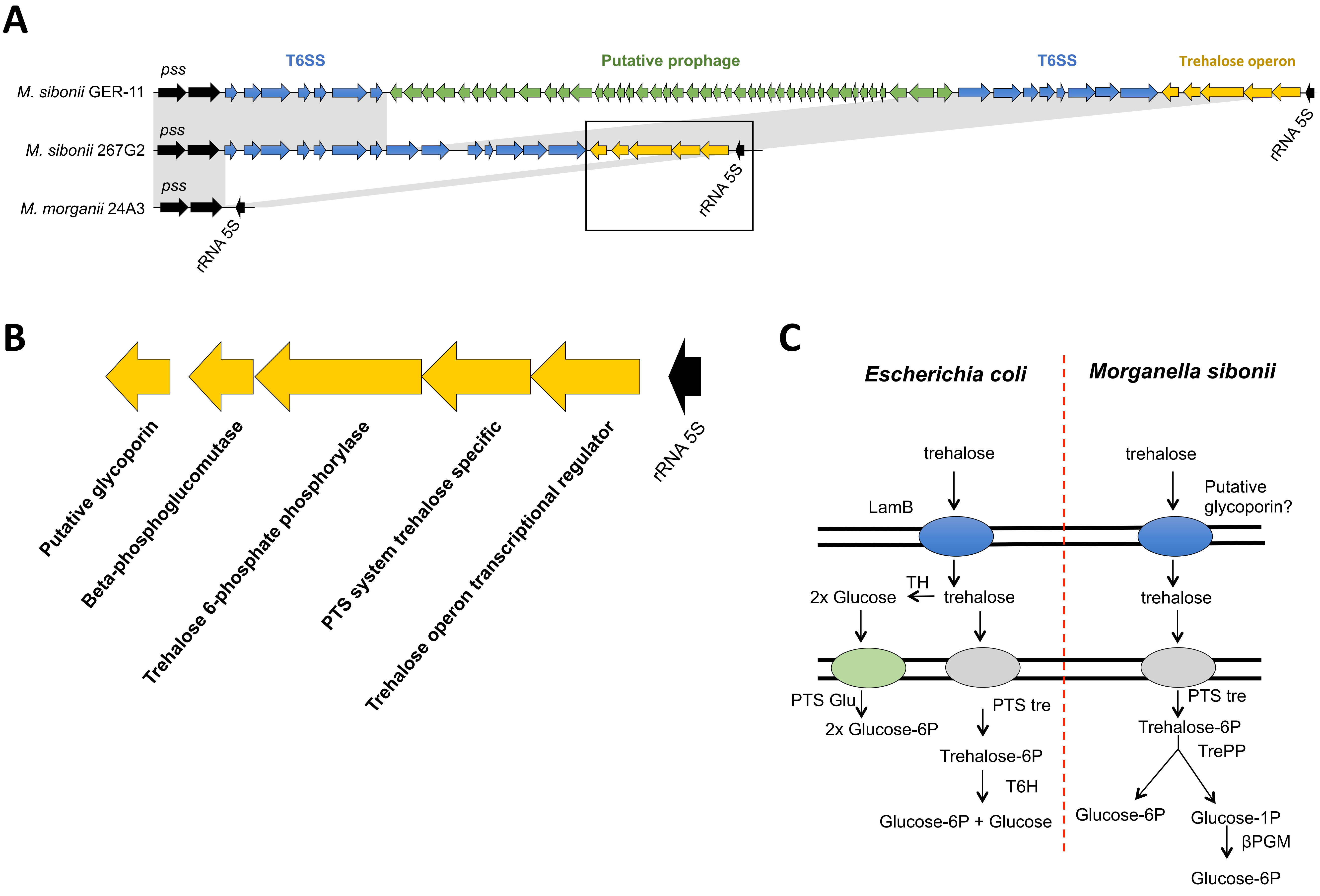

### Supplementary Figure S9

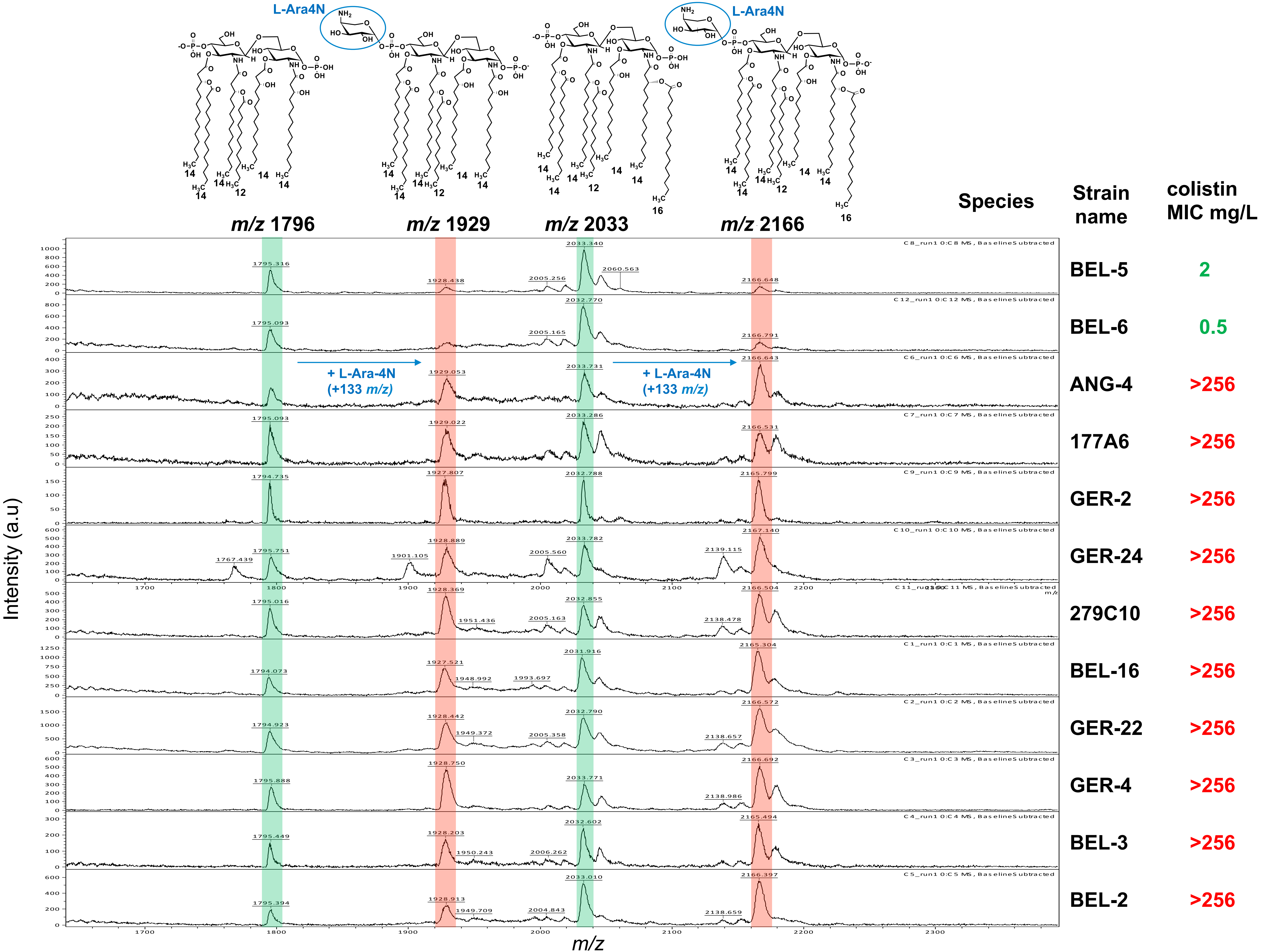

### Supplementary Figure S10

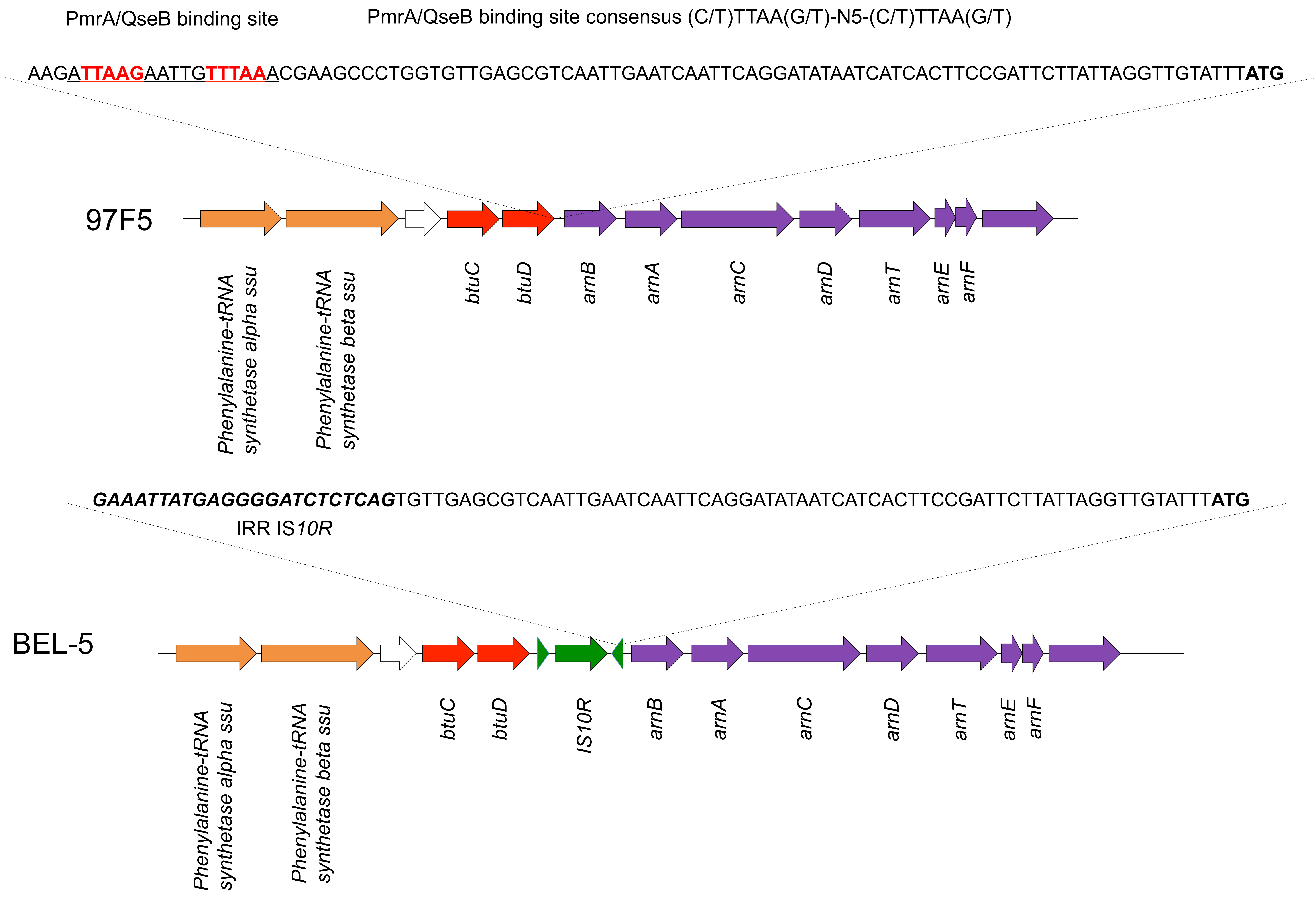

### Supplementary Figure S11

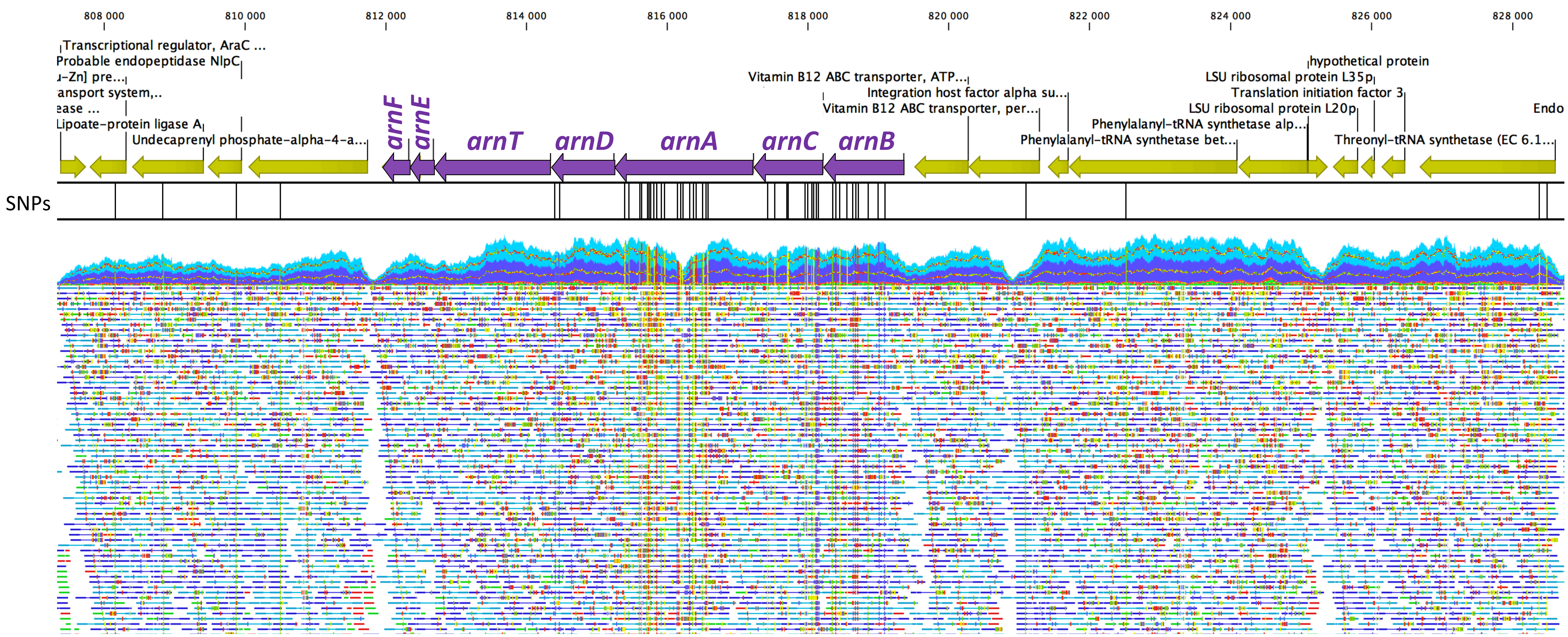
