## Supplementary Figure S6 for "Comparative genomics of carbapenemase-producing *Morganella spp*"

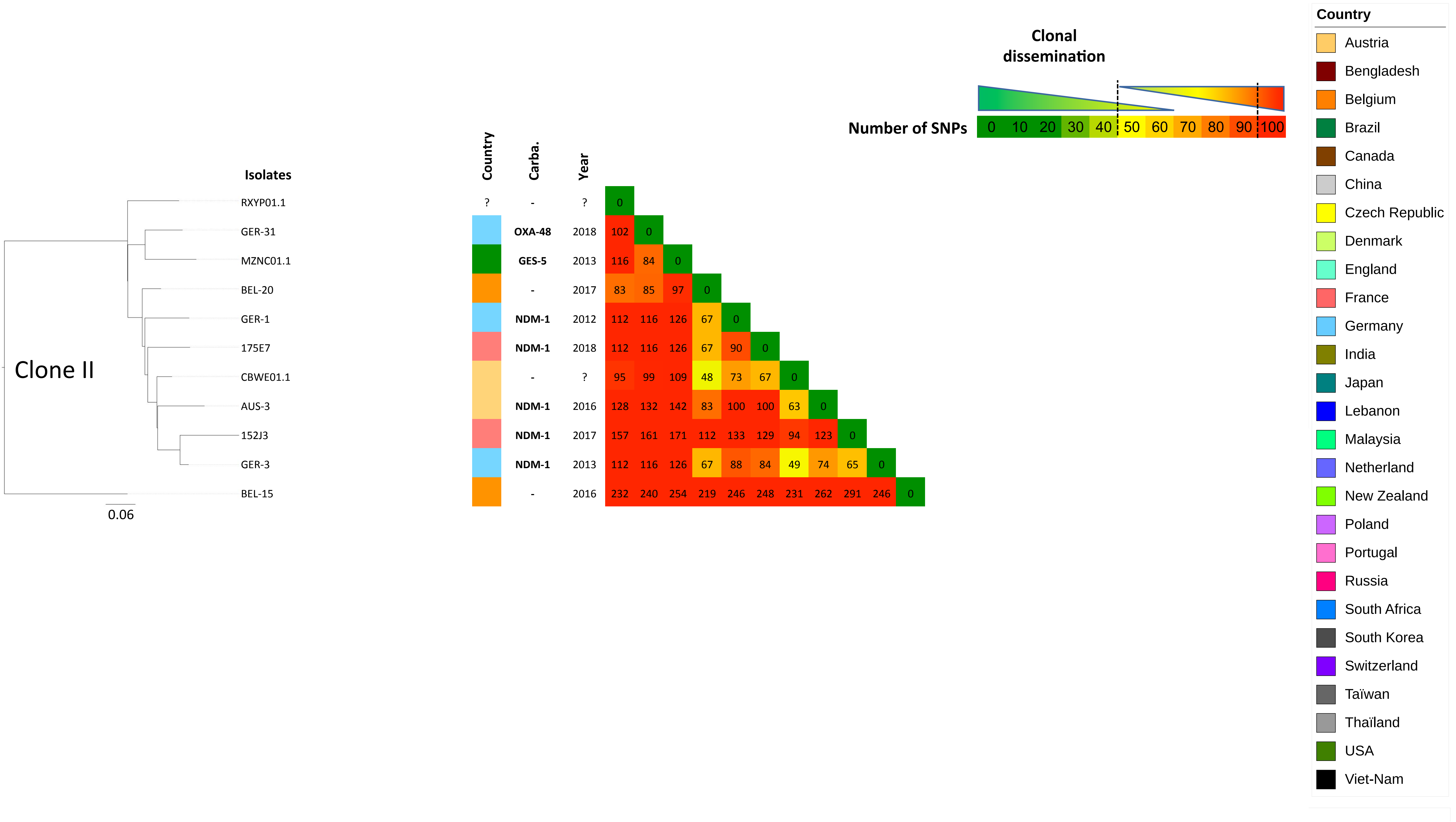

Clonal dissemination
Number of SNPs
0 10 20 30 40 50 60 70 80 90 100
Country
Austria
Bengladesh
Belgium
Brazil
Canada
China
Czech Republic
Denmark
England
France
Germany
India
Japan
Lebanon
Malaysia
Netherland
New Zealand
Poland
Portugal
Russia
South Africa
South Korea
Switzerland
Taïwan
Thaïland
USA
Viet-Nam
Isolates
Country
Carba.
Year
Clone II
0.06
RXYP01.1 ? - ? 0
GER-31 OXA-48 2018 102 0
MZNC01.1 GES-5 2013 116 84 0
BEL-20 - 2017 83 85 97 0
GER-1 NDM-1 2012 112 116 126 67 0
175E7 NDM-1 2018 112 116 126 67 90 0
CBWE01.1 - ? 95 99 109 48 73 67 0
AUS-3 NDM-1 2016 128 132 142 83 100 100 63 0
152J3 NDM-1 2017 157 161 171 112 133 129 94 123 0
GER-3 NDM-1 2013 112 116 126 67 88 84 49 74 65 0
BEL-15 - 2016 232 240 254 219 246 248 231 262 291 246 0

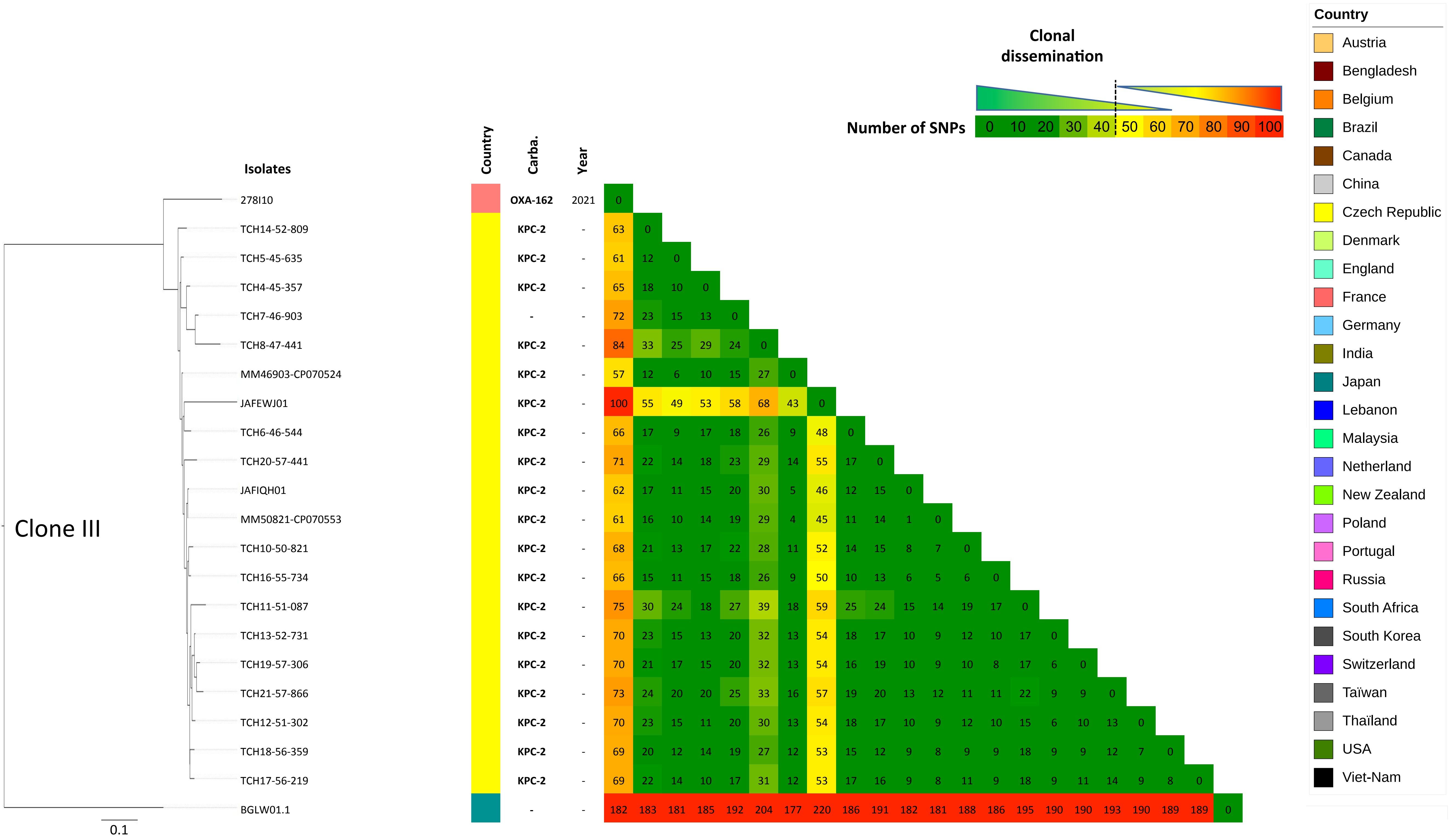

Clonal dissemination
Number of SNPs
0 10 20 30 40 50 60 70 80 90 100
Country
Austria
Bengladesh
Belgium
Brazil
Canada
China
Czech Republic
Denmark
England
France
Germany
India
Japan
Lebanon
Malaysia
Netherland
New Zealand
Poland
Portugal
Russia
South Africa
South Korea
Switzerland
Taïwan
Thaïland
USA
Viet-Nam
Isolates
Country
Carba.
Year
Clone III
0.1
278I10 OXA-162 2021 0
TCH14-52-809 KPC-2 - 63 0
TCH5-45-635 KPC-2 - 61 12 0
TCH4-45-357 KPC-2 - 65 18 10 0
TCH7-46-903 - - 72 23 15 13 0
TCH8-47-441 KPC-2 - 84 33 25 29 24 0
MM46903-CP070524 KPC-2 - 57 12 6 10 15 27 0
JAFEWJ01 KPC-2 - 100 55 49 53 58 68 43 0
TCH6-46-544 KPC-2 - 66 17 9 17 18 26 9 48 0
TCH20-57-441 KPC-2 - 71 22 14 18 23 29 14 55 17 0
JAFIQH01 KPC-2 - 62 17 11 15 20 30 5 46 12 15 0
MM50821-CP070553 KPC-2 - 61 16 10 14 19 29 4 45 11 14 1 0
TCH10-50-821 KPC-2 - 68 21 13 17 22 28 11 52 14 15 8 7 0
TCH16-55-734 KPC-2 - 66 15 11 15 18 26 9 50 10 13 6 5 6 0
TCH11-51-087 KPC-2 - 75 30 24 18 27 39 18 59 25 24 15 14 19 17 0
TCH13-52-731 KPC-2 - 70 23 15 13 20 32 13 54 18 17 10 9 12 10 17 0
TCH19-57-306 KPC-2 - 70 21 17 15 20 32 13 54 16 19 10 9 10 8 17 6 0
TCH21-57-866 KPC-2 - 73 24 20 20 25 33 16 57 19 20 13 12 11 11 22 9 9 0
TCH12-51-302 KPC-2 - 70 23 15 11 20 30 13 54 18 17 10 9 12 10 15 6 10 13 0
TCH18-56-359 KPC-2 - 69 20 12 14 19 27 12 53 15 12 9 8 9 9 18 9 9 12 7 0
TCH17-56-219 KPC-2 - 69 22 14 10 17 31 12 53 17 16 9 8 11 9 18 9 11 14 9 8 0
BGLW01.1 - - 182 183 181 185 192 204 177 220 186 191 182 181 188 186 195 190 190 193 190 189 189 0

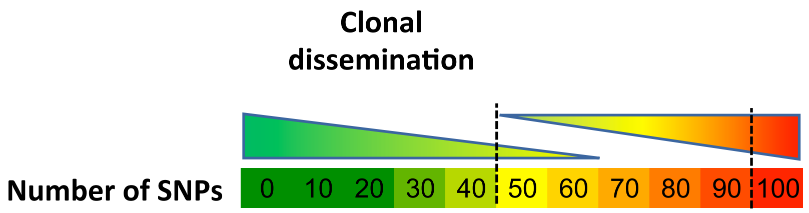


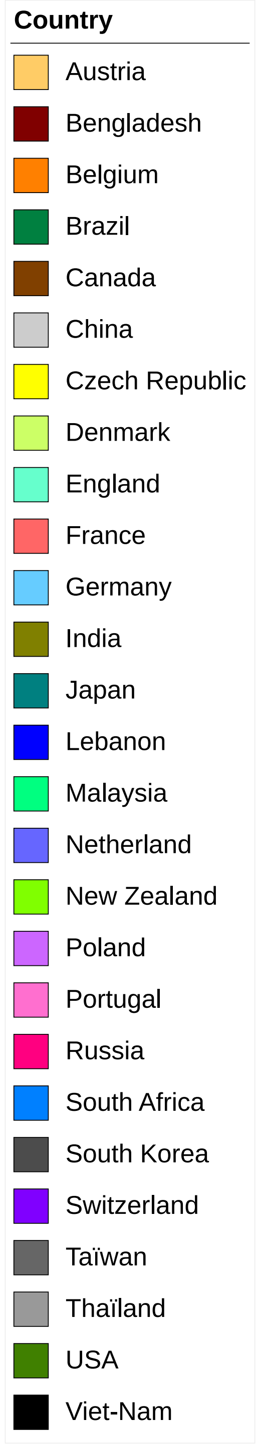


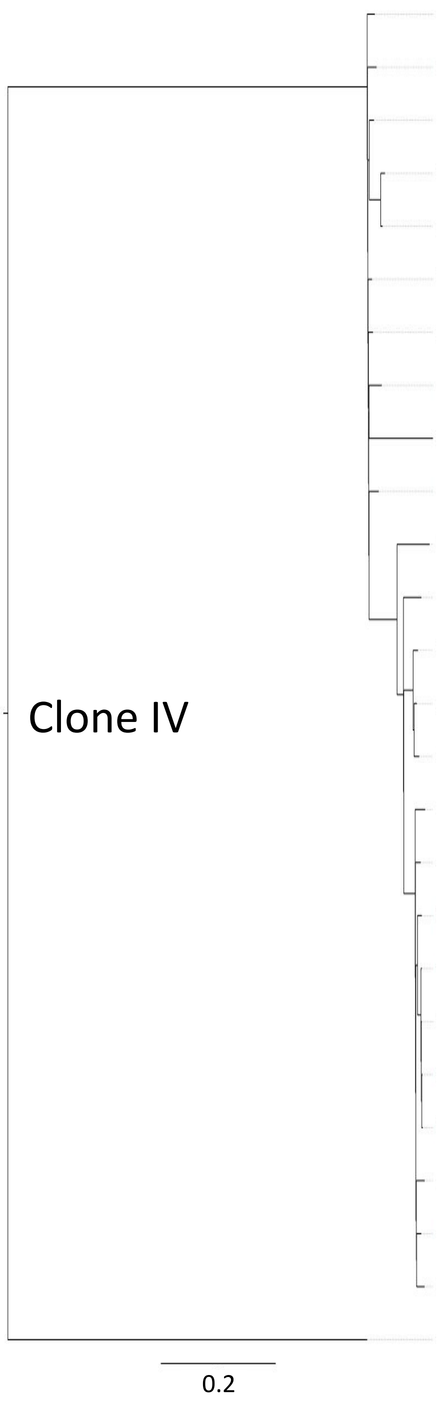


| Isolates | Country | Carba. | Year | | | | | | | | | | | | | | | | | | | | | | | | | | |
|---|---|---|---|---|---|---|---|---|---|---|---|---|---|---|---|---|---|---|---|---|---|---|---|---|---|---|---|---|---|
| GN28_CP026651 | | - | 2015 | 0 | | | | | | | | | | | | | | | | | | | | | | | | | |
| VKWT01 | | - | 2014 | 97 | 0 | | | | | | | | | | | | | | | | | | | | | | | | |
| GER-2 | | - | 2013 | 82 | 89 | 0 | | | | | | | | | | | | | | | | | | | | | | | |
| GER-27 | | NDM-1 | 2018 | 145 | 146 | 117 | 0 | | | | | | | | | | | | | | | | | | | | | | |
| GER-4 | | NDM-1 | 2014 | 131 | 132 | 103 | 36 | 0 | | | | | | | | | | | | | | | | | | | | | |
| BEL-16 | | - | 2016 | 69 | 76 | 59 | 122 | 108 | 0 | | | | | | | | | | | | | | | | | | | | |
| BEL-3 | | OXA-48 | 2014 | 74 | 79 | 64 | 127 | 113 | 49 | 0 | | | | | | | | | | | | | | | | | | | |
| VDHP01 | | - | - | 120 | 127 | 110 | 173 | 159 | 97 | 100 | 0 | | | | | | | | | | | | | | | | | | |
| CCFB01.1 | | NDM-1 | - | 386 | 393 | 374 | 439 | 425 | 363 | 366 | 412 | 0 | | | | | | | | | | | | | | | | | |
| ANG-4 | | NDM-1 | 2016 | 101 | 110 | 93 | 156 | 142 | 78 | 83 | 129 | 395 | 0 | | | | | | | | | | | | | | | | |
| RQIJ01.1 | | - | 2017 | 370 | 377 | 360 | 423 | 409 | 347 | 350 | 396 | 656 | 379 | 0 | | | | | | | | | | | | | | | |
| O86E1 | | - | 2021 | 338 | 345 | 328 | 391 | 377 | 315 | 318 | 364 | 624 | 347 | 316 | 0 | | | | | | | | | | | | | | |
| POL1 | | NDM-1 | 2018 | 321 | 328 | 311 | 374 | 360 | 298 | 299 | 347 | 606 | 330 | 299 | 185 | 0 | | | | | | | | | | | | | |
| GER-15 | | OXA-48 | 2016 | 315 | 320 | 305 | 368 | 354 | 292 | 293 | 341 | 601 | 324 | 293 | 179 | 48 | 0 | | | | | | | | | | | | |
| WCIT01 | | - | - | 299 | 306 | 289 | 352 | 338 | 276 | 279 | 325 | 587 | 308 | 293 | 185 | 64 | 46 | 0 | | | | | | | | | | | |
| NXKH01.1 | | - | 2013 | 357 | 364 | 347 | 410 | 396 | 334 | 339 | 385 | 645 | 368 | 339 | 227 | 204 | 198 | 204 | 0 | | | | | | | | | | |
| FDAARGOS_63_CP026046 | | - | 2013 | 334 | 341 | 324 | 387 | 373 | 311 | 316 | 362 | 622 | 345 | 314 | 198 | 177 | 171 | 177 | 89 | 0 | | | | | | | | | |
| TCH15-54-294 | | OXA-48 | - | 340 | 345 | 330 | 393 | 379 | 316 | 318 | 366 | 624 | 347 | 318 | 204 | 181 | 173 | 181 | 95 | 66 | 0 | | | | | | | | |
| TCH2-37-590 | | - | - | 340 | 346 | 331 | 394 | 380 | 317 | 319 | 367 | 625 | 348 | 319 | 205 | 182 | 174 | 182 | 96 | 67 | 47 | 0 | | | | | | | |
| MM48659-CP070537 | | KPC-2 | | 334 | 342 | 325 | 388 | 374 | 311 | 315 | 361 | 619 | 344 | 313 | 199 | 176 | 170 | 176 | 90 | 61 | 49 | 10 | 0 | | | | | | |
| TCH3-42-744 | | KPC-2 | - | 342 | 348 | 333 | 396 | 382 | 319 | 321 | 369 | 627 | 350 | 321 | 205 | 182 | 176 | 184 | 98 | 69 | 51 | 14 | 10 | 0 | | | | | |
| TCH9-48-659 | | - | - | 344 | 350 | 335 | 398 | 384 | 321 | 323 | 371 | 628 | 352 | 323 | 209 | 182 | 178 | 186 | 100 | 71 | 53 | 16 | 10 | 10 | 0 | | | | |
| 249E6 | | NDM-1 | 2020 | 353 | 360 | 341 | 404 | 390 | 330 | 333 | 379 | 639 | 362 | 333 | 219 | 196 | 190 | 196 | 108 | 81 | 85 | 86 | 80 | 88 | 90 | 0 | | | |
| 156C10 | | NDM-1 | 2017 | 333 | 338 | 325 | 388 | 374 | 312 | 313 | 363 | 623 | 346 | 315 | 201 | 176 | 170 | 178 | 90 | 61 | 65 | 63 | 61 | 67 | 69 | 76 | 0 | | |
| BEL-22 | | - | 2017 | 354 | 359 | 344 | 407 | 393 | 331 | 334 | 382 | 641 | 363 | 334 | 220 | 193 | 189 | 197 | 109 | 80 | 80 | 83 | 81 | 78 | 78 | 95 | 73 | 0 | |
| BFCJ01.1 | | NDM-1 | 2016 | 1995 | 2004 | 1981 | 2050 | 2036 | 1974 | 1979 | 2025 | 2198 | 2006 | 2263 | 2231 | 2214 | 2208 | 2192 | 2250 | 2227 | 2233 | 2234 | 2228 | 2234 | 2238 | 2246 | 2228 | 2247 | 0 |

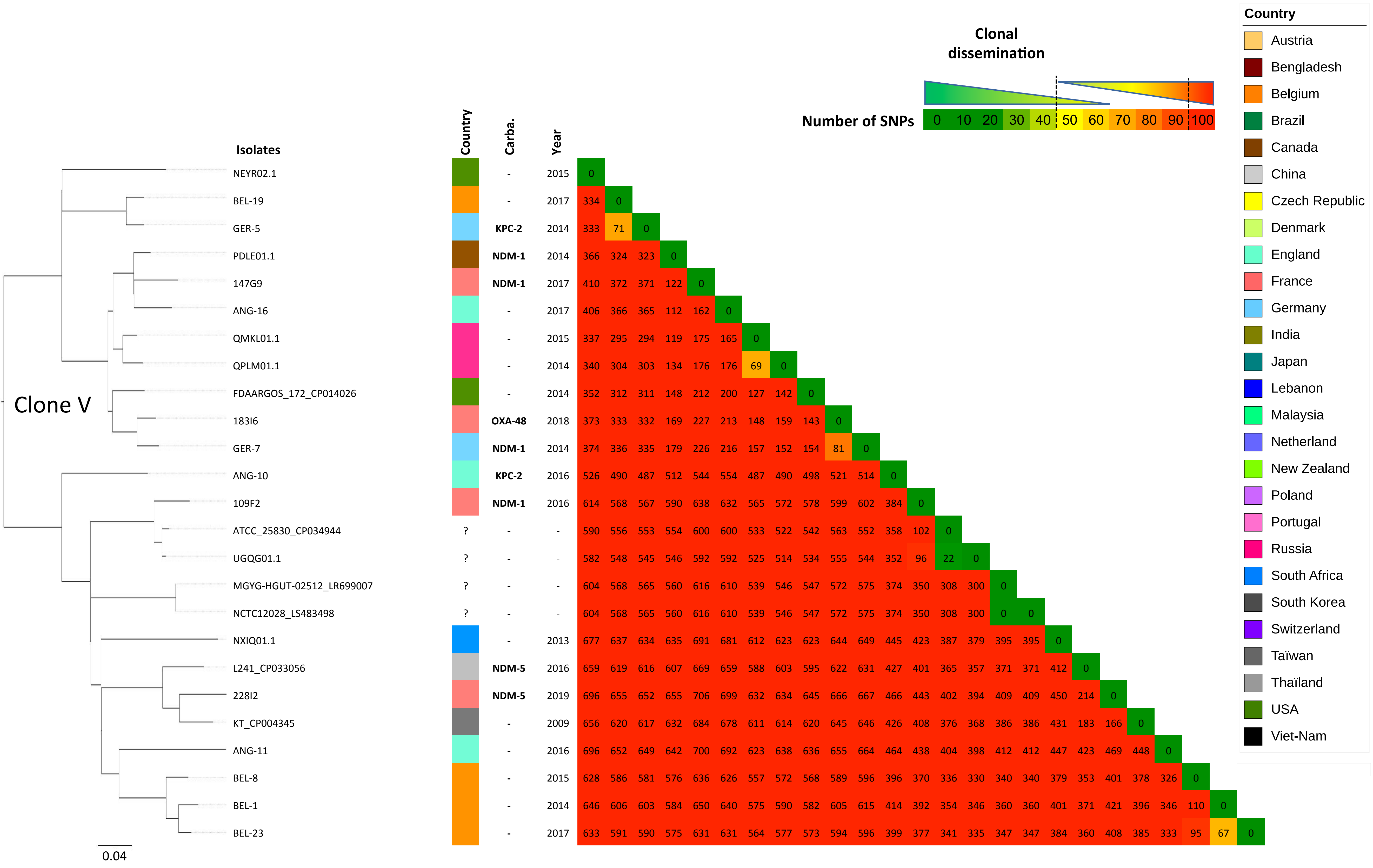

Clonal dissemination
Number of SNPs
0 10 20 30 40 50 60 70 80 90 100
Country
Austria
Bengladesh
Belgium
Brazil
Canada
China
Czech Republic
Denmark
England
France
Germany
India
Japan
Lebanon
Malaysia
Netherland
New Zealand
Poland
Portugal
Russia
South Africa
South Korea
Switzerland
Taïwan
Thaïland
USA
Viet-Nam
Clone V
0.04
Isolates
Country
Carba.
Year
NEYR02.1 - 2015 0
BEL-19 - 2017 334 0
GER-5 KPC-2 2014 333 71 0
PDLE01.1 NDM-1 2014 366 324 323 0
147G9 NDM-1 2017 410 372 371 122 0
ANG-16 - 2017 406 366 365 112 162 0
QMKL01.1 - 2015 337 295 294 119 175 165 0
QPLM01.1 - 2014 340 304 303 134 176 176 69 0
FDAARGOS_172_CP014026 - 2014 352 312 311 148 212 200 127 142 0
183I6 OXA-48 2018 373 333 332 169 227 213 148 159 143 0
GER-7 NDM-1 2014 374 336 335 179 226 216 157 152 154 81 0
ANG-10 KPC-2 2016 526 490 487 512 544 554 487 490 498 521 514 0
109F2 NDM-1 2016 614 568 567 590 638 632 565 572 578 599 602 384 0
ATCC_25830_CP034944 ? - - 590 556 553 554 600 600 533 522 542 563 552 358 102 0
UGQG01.1 ? - - 582 548 545 546 592 592 525 514 534 555 544 352 96 22 0
MGYG-HGUT-02512_LR699007 ? - - 604 568 565 560 616 610 539 546 547 572 575 374 350 308 300 0
NCTC12028_LS483498 ? - - 604 568 565 560 616 610 539 546 547 572 575 374 350 308 300 0 0
NXIQ01.1 - 2013 677 637 634 635 691 681 612 623 623 644 649 445 423 387 379 395 395 0
L241_CP033056 NDM-5 2016 659 619 616 607 669 659 588 603 595 622 631 427 401 365 357 371 371 412 0
228I2 NDM-5 2019 696 655 652 655 706 699 632 634 645 666 667 466 443 402 394 409 409 450 214 0
KT_CP004345 - 2009 656 620 617 632 684 678 611 614 620 645 646 426 408 376 368 386 386 431 183 166 0
ANG-11 - 2016 696 652 649 642 700 692 623 638 636 655 664 464 438 404 398 412 412 447 423 469 448 0
BEL-8 - 2015 628 586 581 576 636 626 557 572 568 589 596 396 370 336 330 340 340 379 353 401 378 326 0
BEL-1 - 2014 646 606 603 584 650 640 575 590 582 605 615 414 392 354 346 360 360 401 371 421 396 346 110 0
BEL-23 - 2017 633 591 590 575 631 631 564 577 573 594 596 399 377 341 335 347 347 384 360 408 385 333 95 67 0
